# Systematic Generation of Mutations at Topoisomerase II Cleavage Sites Enables Cancer Adaptation to Doxorubicin Therapy

**DOI:** 10.64898/2026.08.18.745411

**Authors:** Valid Gahramanov, Anjana Edathil Kadangodan, Riham Barazi, Arkadi Hesin, Julia Yaglom, Vadim E. Levit, Yaakov Maman, Michael Sherman

## Abstract

Acquired resistance to chemotherapy remains a major cause of treatment failure. Here, we investigate the early events underlying the development of resistance to doxorubicin (Dox), one of the most widely used anticancer agents, during repeated drug exposure. Dox intercalates into DNA at sites influenced by chromatin structure and inhibits the DNA religation step of topoisomerase II (Top2), thereby inducing double-strand DNA breaks (DSBs). Here, we show that, in the subpopulation of cancer cells that survive initial Dox treatment, repair of Dox-induced DSBs leads to the accumulation of recurrent mutations at Top2 cleavage sites. These mutations prevent the formation of Dox-induced DSBs at the same loci upon subsequent exposures, effectively protecting the genome from Top2-mediated DNA damage and promoting the development of drug resistance. Importantly, the loss of Top2-dependent DNA cleavage and relaxation caused by these mutations increases the reliance of adapted cells on topoisomerase I (Top1), thereby sensitizing them to Top1 inhibitors. Together, our findings reveal a previously unrecognized mechanism of resistance based on the active generation of highly adaptive mutations, providing new insight into cancer evolution under therapeutic pressure.

## INTRODUCTION

One of the primary causes of treatment failure is the development of drug resistance. In experimental models designed to select drug-resistant mutants in cancer cells, successive passages with escalating drug doses are often used. Some such experiments, in which cell barcoding has been used, obtained a set of resistant clones harboring identical barcodes, indicating that the pre-existing resistance mutations were largely selected upon drug exposures [1–4] [5,6]. In contrast, other selected resistant clones carried unique barcodes across parallels, suggesting that their resistance arises from mutations generated in the process of drug treatment. However, such clones were not further investigated in these studies [5,7,8].

Furthermore, since the initial cell population in experiments that investigate development of drug resistance may originate from a single genetically homogeneous clone, the entire process cannot solely involve the selection of preexisting mutations but must involve the generation of drug-resistant mutations *de novo*. Indeed, previous studies uncovered the activation of mutation-generating mechanisms upon exposure of cancer cells to various drugs [9–13]. Furthermore, these mechanisms, e.g., cytosine deamination by Apobec3A, were functionally involved in the development of resistance to multiple drugs [14–17]. However, the mechanistic role of these drug-induced mutations in the development of drug resistance has not been explored.

In a recent study, we examined the development of resistance to a common drug, irinotecan, which inhibits topoisomerase 1 (Top1) and consequently induces the formation of DNA breaks, including double-strand breaks (DSB), ultimately resulting in cell death [18]. In this study, we unveiled an unanticipated mechanism of irinotecan resistance that is independent of mutations in the drug target, MDR activity, or the expression of genes traditionally associated with the resistance. Instead, resistance emerged through gradual accumulation of recurrent mutations in Top1-dependent DNA-break sites. These mutations reduce the likelihood of Top1-dependent DNA breaks at these sites upon subsequent irinotecan exposures, thereby progressively providing resistance [18,19].

Here, we explore the development of resistance to Doxorubicin (Dox), a widely used Top2 inhibitor, with broad efficacy against a wide array of cancer types [20–23]. Under normal conditions, Top2 relieves torsional stress in DNA by transiently binding and introducing reversible DSBs, allowing DNA stress relaxation. In the presence of Dox, the re-ligation step is blocked, leading to the accumulation of DSBs [24–26]. Previously, it was shown that multiple mechanisms, such as overexpression of MDR pumps, mutations in Top2, or activation of pathways that affect cell death, play an important role in Dox resistance [27–31]. Here, we uncovered a novel major mechanism, which involves a gradual accumulation of mutations in Top2 cleavage sites. These mutations progressively diminish the ability of Top2 to generate DSBs at Dox-induced break sites (DIB), leading to a reduction in Dox toxicity.

## RESULTS

### Experimental design - dose escalation Dox treatments

As a model to study the development of resistance to doxorubicin (Dox), we used the colon cancer cell line HCT116. To maximize genetic homogeneity, cells were single-cell cloned and independent clones were isolated (Fig. 1a). This approach ensured that resistance-associated mutations arising during treatment were generated *de novo* rather than originating from pre-existing subclones. Although some genetic divergence may occur during clonal expansion, it is unlikely to significantly affect our analysis, as the mutation patterns observed were drug-specific (see comparison between Dox- and irinotecan-induced mutations, Fig. S5a,b).

**Figure 1.**
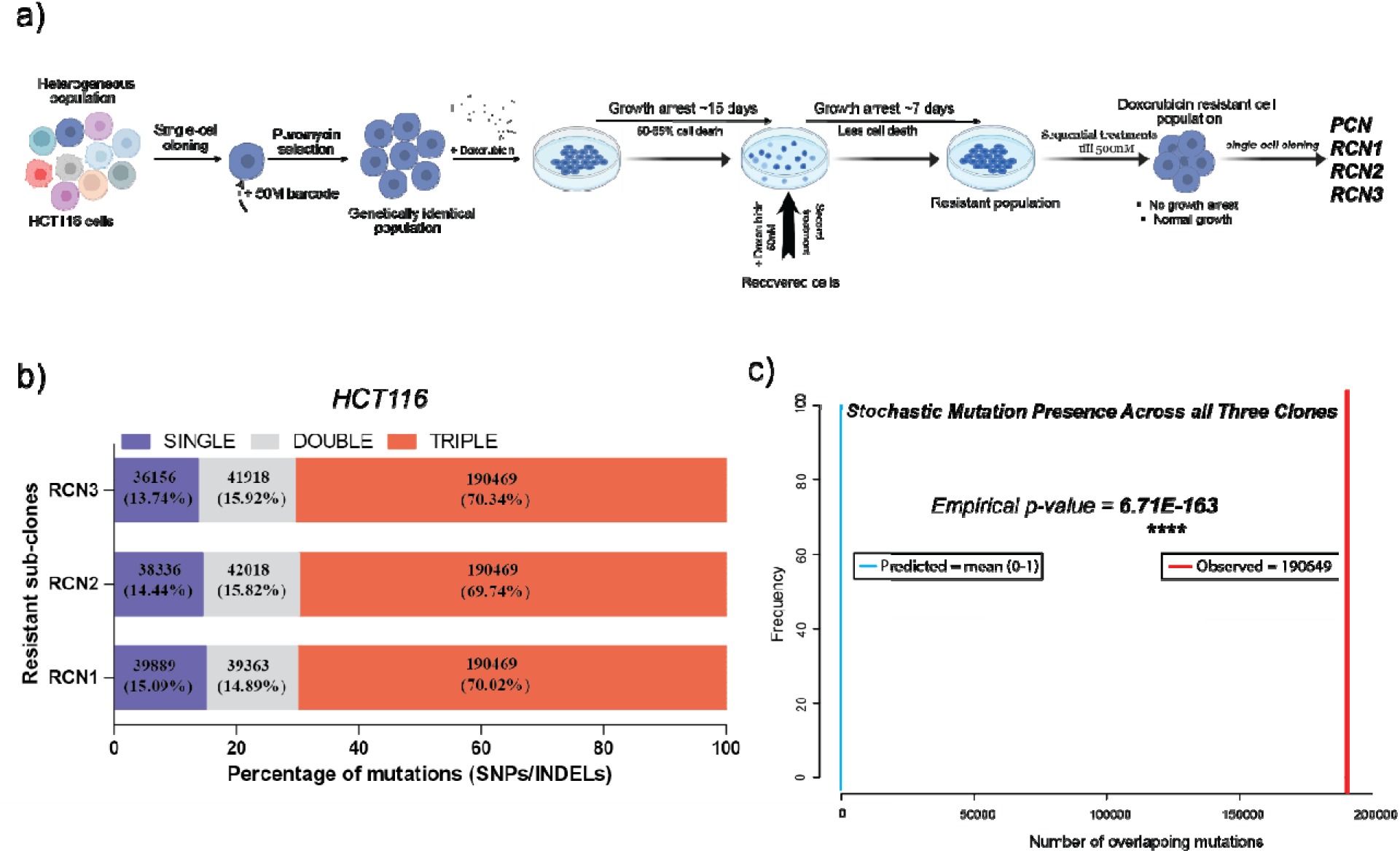
Adaptation to Dox. (a) Schematic overview of the experimental design for the generation of Dox-resistant HCT116 cells (RCN1, RCN2, RCN3). (b) Quantification of overlapping mutations among the Dox-resistant clones that emerged during the adaptation process. The stack bar shows the number of overlapping mutations among resistant clones. These mutations represent three categories: mutations that are unique to each clone and not found in other clones (blue), mutations that are identical in two out of three clones (gray), and mutations that are identical in all three clones (red). (c) The plot illustrates th statistical comparison between the observed and expected number of common mutations shared among three clones. The red line represents the actual number of overlapping mutations observed in all three clones. The sky-blue bars depict the distribution of overlap counts obtained from 100 permutations of randomly generated genomic mutation sites. Statistical significance (**** p < 0.00001) was assessed using a Monte Carlo permutation test, evaluating whether the observed overlap exceeds what would b expected by chance. The y-axis indicates the number of permutations contributing to the null distribution.

To enable lineage tracing during treatment, cells from the expanded parental clone were uniquely DNA-barcoded prior to Dox exposure. Thus, cells carrying distinct barcodes at the end of the selection process represent independent lineages that survived the full series of treatments.

Cells were initially exposed to 50 nM Dox, resulting in the death of approximately 40–50% of the population. The surviving cells entered a growth-arrested, senescence-like state characterized by enlarged morphology, vacuolization, and increased senescence-associated β-galactosidase activity. This pseudo-senescent phase lasted approximately 15 days and was followed by resumption of proliferation. With repeated cycles of 50 nM Dox treatment, the duration of growth arrest progressively decreased. By the fourth cycle, minimal cell death or growth inhibition was observed, indicating effective adaptation to this dose (Figs. 1a, S1, S2). Cells were subsequently adapted to progressively higher concentrations, reaching 500 nM Dox.

**Figure 2.**
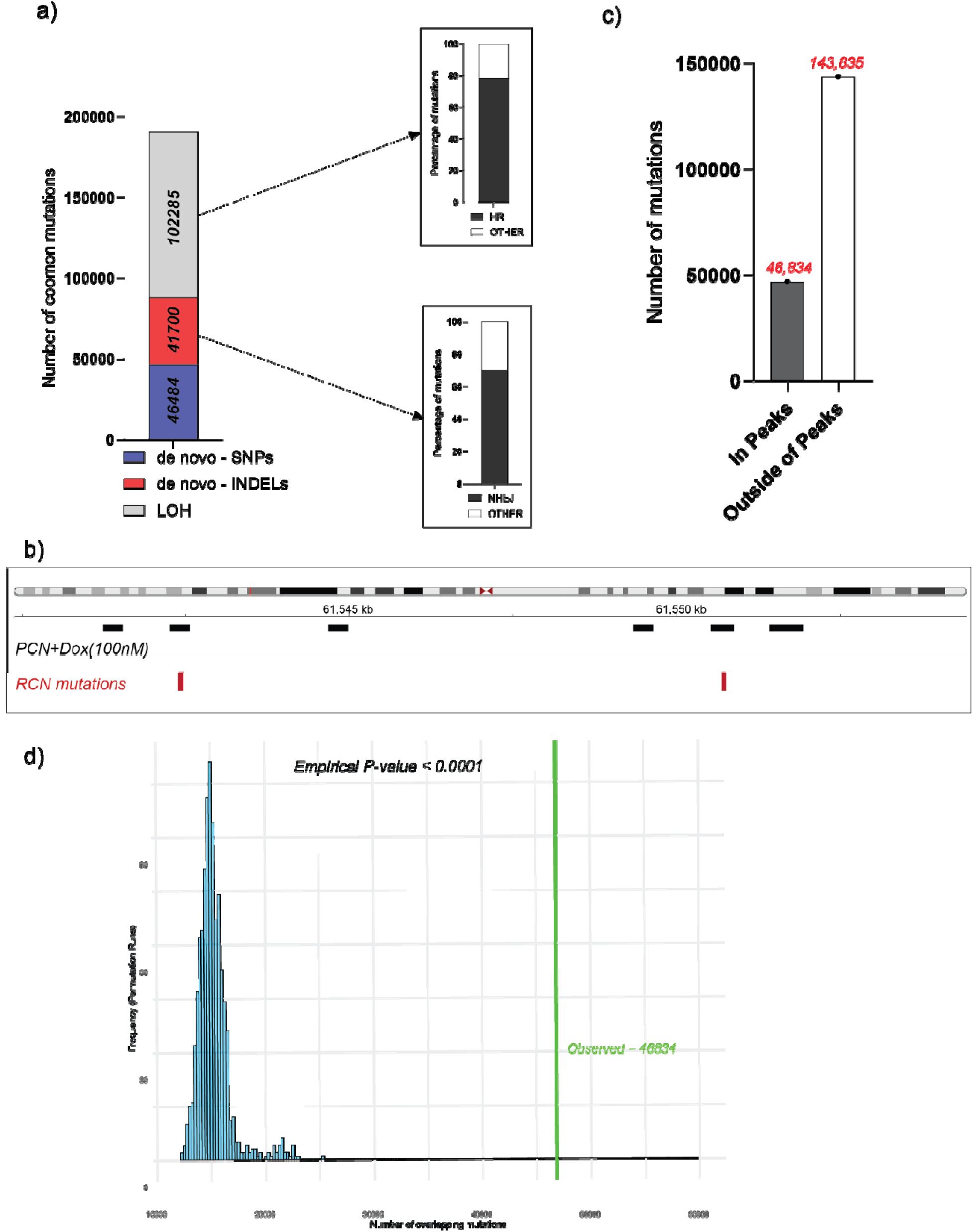
DNA repair through HR and NHEJ mechanisms can lead to characteristic mutation signatures. (a) Categorical distribution of common mutations classified as *de novo* SNPs (Table S3c), *de novo* INDELs, and loss of heterozygosity (LOH) mutations. The right panels show bar graphs representing the proportion of LOH attributed to homologous recombination (HR) and *de novo* InDels attributed to non-homologous end-joining (NHEJ) repair mechanisms. (b) Co-localization of END-seq peaks (recurrent DNA break sites) (black) in PCN cells treated with 100□nM Dox and the common mutations (red) shared by all three RCN clones, is shown in a representative genomic region (chr1: 61,540,000 – 61,555,500). (c) Bar graph depicting a high proportion of colocalization of END-seq peaks with common mutations. ‘In peaks’ indicates the number of mutations located within END-seq peaks, while ‘out of peaks’ represents mutations found outside of the peak regions. (d) The permutation test distribution plot illustrates the difference between the observed and expected overlap of common mutations with END-seq peaks. Randomized mutation sites were generated 100 times, and in every iteration, the number of overlaps with END-seq peaks was significantly lower than the observed overlap (46,834 mutations), yielding an empirical p-value of <0.0001. The green line indicates the observed number of mutations that overlapped with the END-seq peaks.

To assess the generality of this adaptive response, we performed parallel experiments in the lung cancer cell line H1975, which differs from HCT116 in both tissue origin and DNA repair capacity and lacks microsatellite instability. Notably, H1975 cells exhibited similar adaptation kinetics (Figs. S1, S2), suggesting that the observed phenomenon is not cell line-specific.

To investigate mechanisms of resistance, cells adapted to 500 nM Dox were recloned. Independent resistant clones derived from distinct barcoded lineages (RCN1, RCN2, and RCN3; RCN, resistant clone) were selected for further analysis (Fig. 1a).

### Limited contribution of transcriptomic changes to resistance development

To identify mechanisms underlying resistance, we first compared the transcriptomes of resistant clones with that of the Dox-sensitive parental clone (PCN). Approximately 1,400 genes were differentially expressed in resistant clones relative to PCN (Table S1), with a high degree of overlap across independently derived resistant clones.

Importantly, we did not detect significant changes in the expression of genes previously implicated in Dox resistance, including *MDR1*, *TOP2A*, *TOP2B*, or key DNA repair genes. Notably, several multidrug resistance transporters were downregulated in resistant clones (Table S1, Fig. S3).

Gene Set Enrichment Analysis (GSEA) revealed limited and inconsistent pathway-level changes: one resistant clone showed enrichment of interferon signaling, whereas another displayed activation of MYC-related pathways (Table S2). These pathways are not directly linked to established mechanisms of Dox resistance. Although some MYC targets may contribute to adaptation, the lack of consistent and mechanistically relevant transcriptional changes across clones suggests that transcriptomic reprogramming is unlikely to be the primary driver of Dox resistance in this system.

### Recurrent, non-random mutations emerge during Dox adaptation

To further investigate the molecular basis of resistance, we performed whole-genome sequencing of resistant clones (RCN1, RCN2, RCN3) and the parental clone (PCN). Genomes of the resistant clones were compared both to the parental clone and to the reference genome (GRCh38). Comparison of RCN and PCN genomes identified approximately 270,000 point mutations and INDELs in each resistant clone that arose during Dox treatment (Fig. 1b). Remarkably, a large fraction of these mutations was shared across independently derived clones: ∼60% of mutations (190,469 total; 51.8% INDELs and 48.2% point mutations) were common to all three clones, with an additional ∼20% shared between two of the three clones (Fig. 1b). This high degree of overlap is striking given the independent origin of these lineages.

A key feature of this system is that all detected mutations were generated *de novo* during the course of repeated drug treatments, as only differences between resistant and parental genomes were considered. Across the dose-escalation protocol, cells underwent approximately 25 cycles of treatment and recovery. Based on the total number of mutations, this corresponds to an average of ∼11,000 mutations per cycle, of which ∼7,600 were identical across all three clones. We therefore assessed the likelihood that such a high proportion of shared mutations could arise by chance. Mathematical modeling, accounting for genome size and total mutation burden, indicated that this probability is effectively zero (P < 3.9 × 10□□²□□□; see Materials and Methods and Supplementary File “Mutation Resonance Probability”). Given that each treatment involved ∼10□ cells, stochastic generation of this level of mutation convergence is practically impossible. These findings strongly suggest that mutation acquisition is not random but instead driven by a specific underlying mechanism. This conclusion is further illustrated by computational simulations (Fig. 1c). Though the probability in the computer simulation was higher than the calculated probability, it was still almost zero.

Consistent with this interpretation, a similarly high fraction of shared mutations was observed in independently derived resistant clones of H1975 cells (Fig. S4), indicating that this phenomenon is not specific to the mismatch repair deficiency (hMLH1 loss) present in HCT116 cells.

Despite the broadly similar mechanisms of action of Dox and the topoisomerase I inhibitor irinotecan—both of which induce DNA breaks—comparison of whole-genome sequencing data revealed distinct mutation distributions across the genome (Figs. S5a,b). Direct comparison of mutation profiles showed only a small subset of overlapping mutations (519 shared sites) (Fig. S6). These results indicate that, although both drugs induce DNA damage, the mechanisms governing the genomic localization of newly generated mutations differ substantially between Dox and irinotecan.

### Mutations arise from repair of Top2-mediated DNA double-strand breaks

To determine the origin of the extensive generation of mutations observed during adaptation to Dox, we hypothesized that these mutations arise from the repair of Top2-mediated DSBs. Consistent with this idea, multiple lines of evidence indicate that the majority of mutations generated during Dox treatment reflect DNA repair events mediated by homologous recombination (HR) or non-homologous end joining (NHEJ).

Analysis of mutation types and repair signatures revealed two dominant patterns corresponding to these pathways. Approximately 55% of mutations resulted in loss of heterozygosity (LOH) (Fig. 2a). In cases where sequencing read depth remained unchanged—excluding deletions—LOH most likely arose through HR-mediated gene conversion, whereby one allele is copied from the homologous chromosome. This mechanism accounted for the majority of LOH events in Dox-resistant clones (Fig. 2a, Table S3a).

In contrast, most de novo insertions and deletions (InDels) bore signatures characteristic of NHEJ (Fig. 2a, Table S3b). In particular, insertion events frequently exhibited short duplications of neighboring regions, consistent with end alignment followed by gap filling through translesion DNA synthesis. These sequence features enable precise localization of the original DNA break sites. Thus, LOH events and InDels serve as complementary markers of DSB repair via HR and NHEJ, respectively.

Importantly, these inferred DSB sites corresponded to known sites of Top2 activity. Accordingly, we compared locations of mutation sites shared across independent resistant clones with published Top2 binding sites, including a dataset generated in HCT116 cells [32]. The overlap between the positions of mutations and Top2 binding sites was dramatically greater than expected by chance based on permutation analysis (Fig. S7), supporting the conclusion that mutations preferentially arise at Top2 cleavage sites. Importantly, comparison of ChIPseq peaks with mutations has a number of caveats, discussed in this section below, and it is not surprising that only a portion of Top2 binding sites (identified by ChIPseq) overlapped with mutations (identified by the genome sequencing).

To further link the Dox-triggered mutations to the sites of the Top2-dependent DNA cleavage in the parental PCN cells, we applied the END-seq approach that directly maps DSBs (see Materials and Methods). To increase the uniformity of cells and facilitate the identification of recurrent break sites, cells were synchronized by the cell cycle inhibitor palbociclib prior to Dox treatment. To a large degree, peaks corresponding to Dox-mediated DSBs were co-localized with the mutations identified in the cells adapted to Dox (see Fig. 2b, as an example). Specifically, a substantial fraction of shared mutations in RCN clones (∼25%) co-localized with DSB peaks in PCN cells, further suggesting that Dox-induced mutations arise at Top2 cleavage sites (Fig. 2c). Permutation analysis indicates that this overlap is unlikely to occur by chance (Fig. 2d). Notably, the difference between the total number of mutations in RCN (300+ thousand) and DSB peaks in PCN (1.2 million) may reflect differences in the experimental setup, see below.

Consistent with the prior report thats CTCF recruits Top2 to facilitate DNA cleavage [32–34], a large fraction of the ENDseq peaks was located at the chromosomal loop anchors occupied by CTCF (Fig. 3a, b), and there were significantly fewer breaks at these locations in RCN cells compared to PCN. Moreover, Dox-induced mutations were highly enriched at CTCF binding sites (Fig. 3c), further suggesting that the decreased breakage in RCN cells was due to the loss of the Top2 binding at these sites.

**Figure 3.**
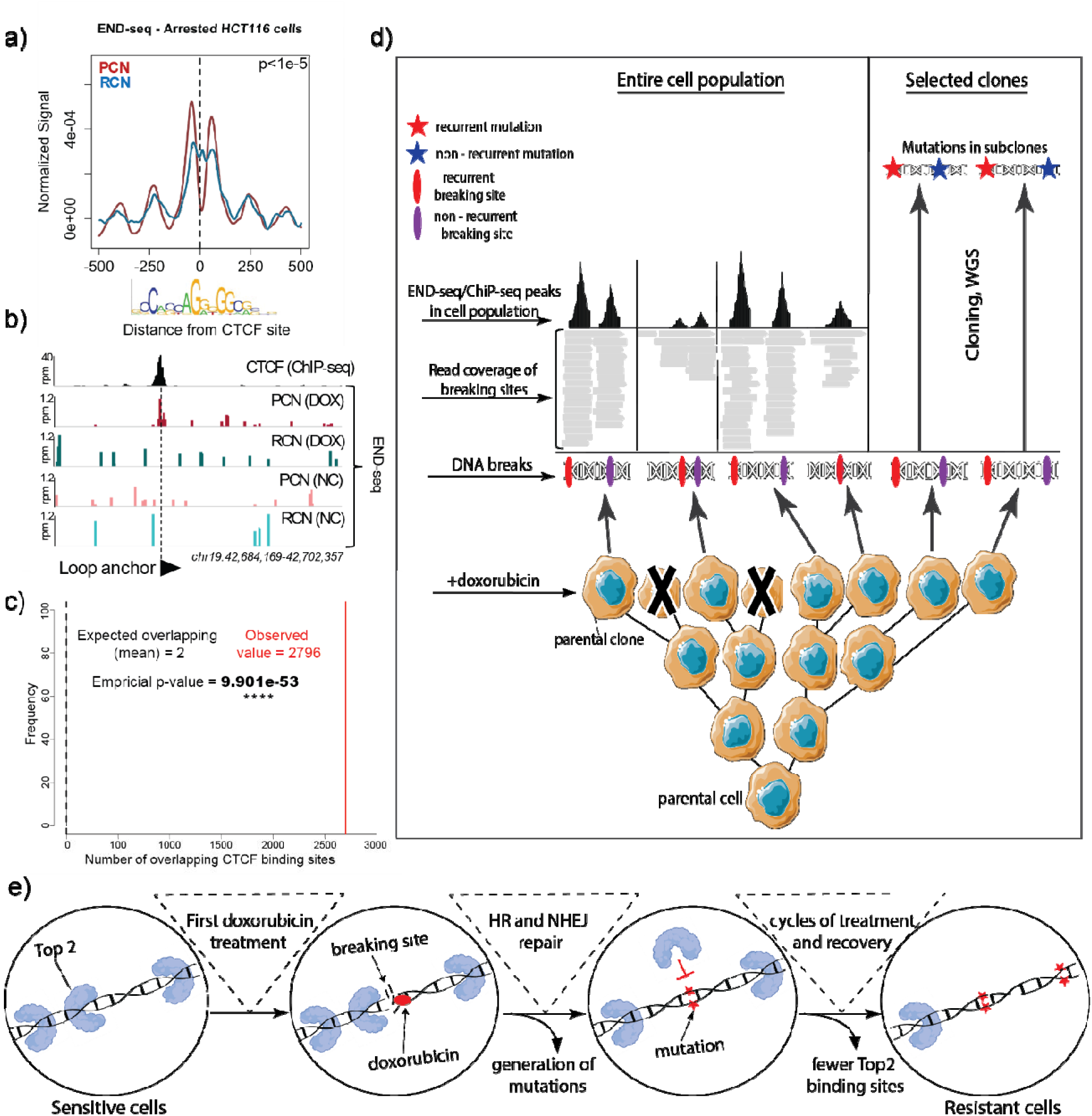
(A) Aggregate plot of Dox-induced DSBs at +/− 500 bp from the CTCF motif in Dox-sensitive (PCN; red), and Dox-resistant (RCN; blue) cells. T-test;p<1e-5 (B) Genome browser snapshot showing (top to bottom): CTCF ChIP-seq signal, and END-seq signal from treated (Dox), and untreated (NC) PCN and RCN cells. (C) The permutation analysis shows the statistical comparison between the observed and expected overlap of common mutation sites with published CTCF-binding regions from the JASPAR database. Statistical significance was assessed using a Monte Carlo permutation test (n = 100 iterations). The p-value (**** p < 0.0001) indicates whether the observed overlap exceeds what would be expected by chance. The x-axis represents the number of overlaps between common mutations and CTCF-binding sites (observed and random), while the y-axis denotes the number of permutation iterations contributing to the null distribution (n = 100). (d) A scheme explaining the caveats in comparison of ENDseq (or ChIPseq) with mutations. Firstly, ENDseq has high background noise and requires a strong filtering, and thus detects only most frequent studies DSB in the cell population. Secondly, mutation analysis, unlike ENDseq, is done with cell cloned after the treatments, and thus detects only mutations in individual clones rather than the entire dox-treated cell population. (e) A model of development of doxorubicin resistance. Repairing DSB following initial exposures generates mutation at these Top2 cleavage sites, which reduces the number of Top2 cleavage events upon further doxorubicin exposures.

A special note regarding the comparison of ENDseq (or ChIPseq) data with mutation data is that the mutation analysis of a cloned population has an incomparably better ability to detect mutations than ENDseq to detect DSBs. Indeed, the genome sequencing identifies all mutations that were present in the cell that formed the clone (Fig. 3d). The ENDseq, however, is dealing with a cell population in which some cells have Top2 bound to a particular site, while in other cells, positions of Top2 could be different. This method can identify only sites where Top2 bound in a significant fraction of cells in the doxorubicin-treated population. In addition, the method has a high noise level. Therefore, the set of ENDseq peaks that represents “clean data” always results from a very significant filtering and corresponds to a sample of most prominent DSB sites in the population.

In addition, mutation analysis identifies mutations in one cell that was expanded to form the RCN clone and thus represents only a subset of mutations in the cell population (Fig. 3d), while ENDseq evaluates DSB in the entire population. Therefore, one can expect only a limited correspondence between ENDseq and mutation analysis. Accordingly, a highly significant overlap between mutations and ENDseq peaks indeed indicated that mutations were generated at the Top2 cleavage sites. Comparison of ChIPseq and mutations data has similar caveats.

The idea that the mutations result from the generation of DSBs fit well with the known mechanism of Dox action via inhibition of the ability of Top2 to re-ligate cleaved DNA. The overlap of mutations in different resistant clones can be attributed to the non-random binding of Top2 to chromatin sites and non-random localization of Dox DNA intercalation sites at the open chromatin [35,36]. The large proportion of shared mutations (about 60% of triple mutations) suggests that the mutations affected the majority of the Dox-induced cleavage sites (Fig. 1b). These findings indicate that repair of the Top2-generated DSBs was the primary cause of the majority of mutations.

### Mechanism of resistance: Adaptation is associated with the reduced ability of Dox to trigger DSB

Based on these findings, we propose that resistance to doxorubicin (Dox) arises through the progressive accumulation of mutations at Top2 cleavage sites. Once mutated during early cycles of Dox exposure, these sites become less permissive to Top2-mediated DNA cleavage. As a result, repeated drug treatments lead to a gradual reduction in functional Top2 cleavage sites across the genome, thereby decreasing the ability of Dox to induce DSBs. In this model, mutations acquired during initial exposures effectively “precondition” the genome, conferring protection against subsequent treatments (Fig. 3e). Notably, this mechanism does not rely on changes in canonical resistance pathways but instead reflects direct alteration of the DNA sequence to reduce susceptibility to Top2-mediated damage.

Independent support for this model comes from analysis of genomic variants in parental and resistant clones relative to the reference genome. The parental PCN genome contained approximately one million variants, including both SNPs and InDels. Strikingly, ∼95% of these variants were absent from the gnomAD database [37], indicating that they largely represent somatic mutations acquired during cancer evolution. We refer to these as “cancer alleles.”

Analysis of the loss of heterozygosity (LOH) mutations upon selection of the resistant RCN clones provides further support for the model in which Top2-directed mutagenesis at the DIB sites leads to Dox resistance. The following considerations provide a framework for understanding the connection between LOH and Dox-resistance. In each genomic position, the alleles within the parental PCN population may exhibit variations: no mutations relative to the reference genome (RG) allele (0/0), mutations in one allele (0/1) indicating heterozygosity, or mutations in both alleles (1/1) (Fig. 3a). Sites where two alleles in PCN had different mutations compared to the RG (1/2) were rare and were excluded from further analysis (Fig. 4a).

**Fig. 4.**
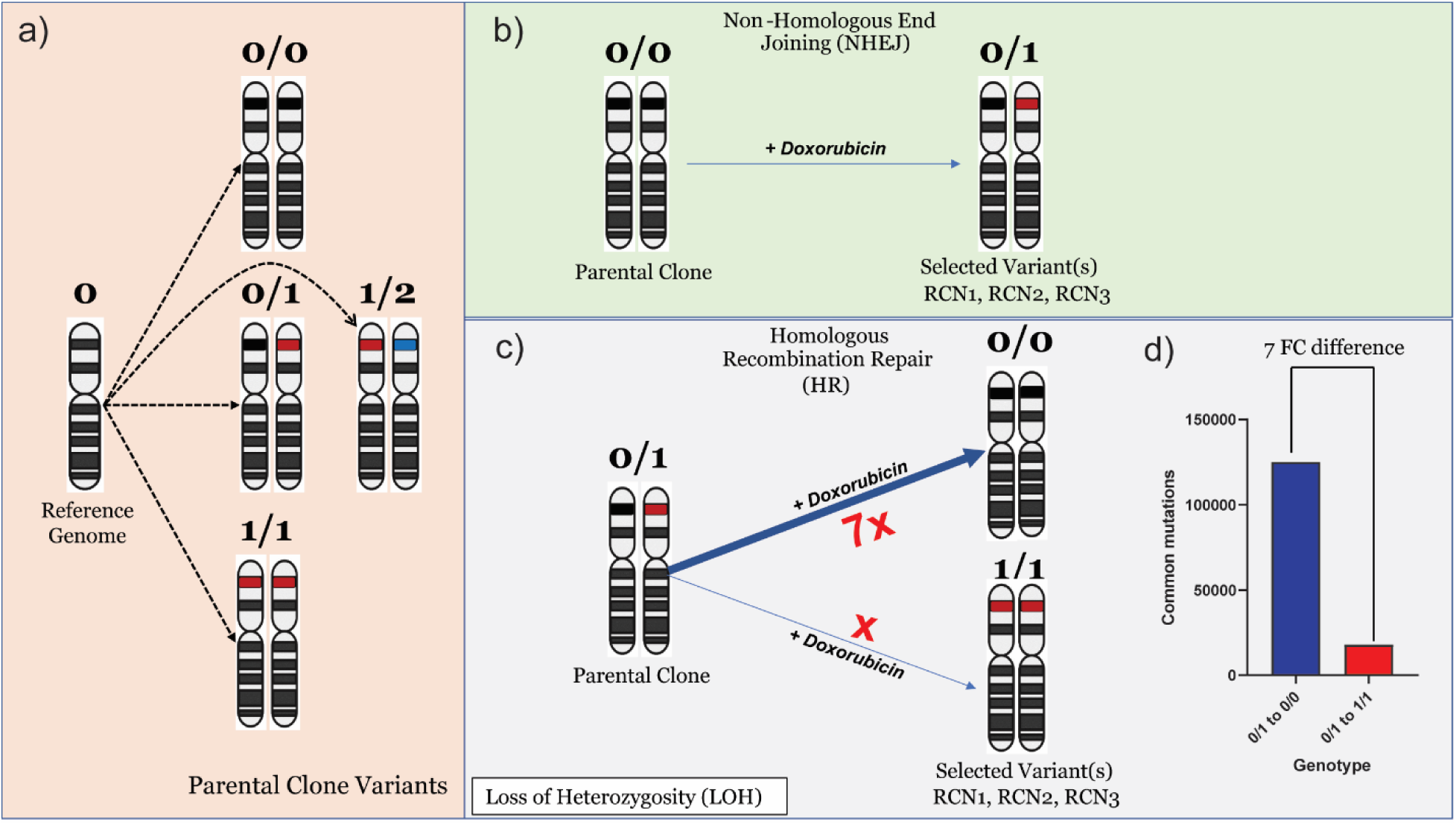
A framework model for analyses of the role of mutations in adaptation to Dox. Schematic representation for (a) allelic variants found in the parental PCN clone compared to the normal reference genome allele, (b) de novo allele variants in RCN clones resulted from the NHEJ repair, (c) allele change upon loss of heterozygosity after HR repair (loss of heterozygosity herein is referred to as the change of mutated heterozygous alleles reversed to homozygous allele in selected variants, such as shift from 0/1 to 0/0 or 0/1 to 1/1). (d) Cancer alleles are more susceptible to breaks compared to reference genome alleles. HR-repair-generated mutations reverted to the reference genome allele seven times more often than to the cancer allele. Therefore, cancer alleles are broken by Top2 seven times more often than reference genome alleles. Accordingly, the reference genome alleles are more resistant to Top2-generated breaks, and upon reverting to these alleles, cells become protected from Top2-mediated DNA cleavage upon consequent exposures to Dox.

Accordingly, mutations detected in RCN clones relative to the parental PCN could arise either as *de novo* InDels (e.g., 0/0 → 0/1) generated by non-homologous end joining (NHEJ) (Fig. 4b) or as loss of heterozygosity (0/1 → 0/0; or 0/1 → 1/1) resulting from homologous recombination (HR) (Fig. 4c). In sites repaired through HR and displaying LOH, accounting for more than 55% of the total mutations, a heterozygous allele may revert either to the reference genome allele (0/1 → 0/0) or the cancer allele observed in the parental clone PCN (0/1 → 1/1). Surprisingly, upon analysis of these mutations, we found that the frequency of 0/1 → 0/0 shifts was 7 times higher than the frequency of 0/1 → 1/1 shifts (Fig. 4c). A shift from 0/1 to 0/0 implies the copying of the RG allele to replace the cancer allele during HR, suggesting that the likelihood of a DSB occurring in cancer alleles is 7 times higher than in the reference genome allele. A similar enrichment pattern was also observed in H1975 cells, where the frequency of 0/1 → 0/0 conversions was 2.3 times higher than that of 0/1 → 1/1 conversions (Fig. S8a, b). Therefore, new alleles acquired during the progression of cancer in the PCN parental clone are significantly more prone to DSB caused by Top2 in the presence of Dox than the corresponding normal reference genome alleles. Thus, it appears that the evolution of cancer cells can lead to the generation of a significant number of additional Dox-induced breaking sites. Accordingly, replacement of these alleles with the normal alleles leads to the elimination of the Top2 sites, which further contributes to the resistance to Top2 cleavage upon the next exposure to Dox. NHEJ-mediated mutations likely contribute to this process as well by disrupting cleavage motifs through insertions and deletions. Together, these results support a mechanism in which selection and repair-driven remodeling of DNA sequence progressively eliminates Top2 cleavage sites, thereby reducing Dox-induced DNA damage and promoting resistance.

### Testing the model

A central prediction of our model is that repeated exposure to Dox should lead to a progressive reduction in Dox-induced DSBs. To test this, we compared DSB levels in PCN and RCN1 cells following treatment with 100nM Dox for 6 hours. DNA damage was quantified by immunostaining for γH2AX (Fig. 5a). As expected, Dox treatment induced a robust increase in γH2AX foci in PCN cells, whereas RCN1 cells exhibited a markedly reduced number of foci under the same conditions (Fig. 5a,b). These results suggest that resistance is associated with a diminished ability of Dox to induce DSBs.

**Figure 5.**
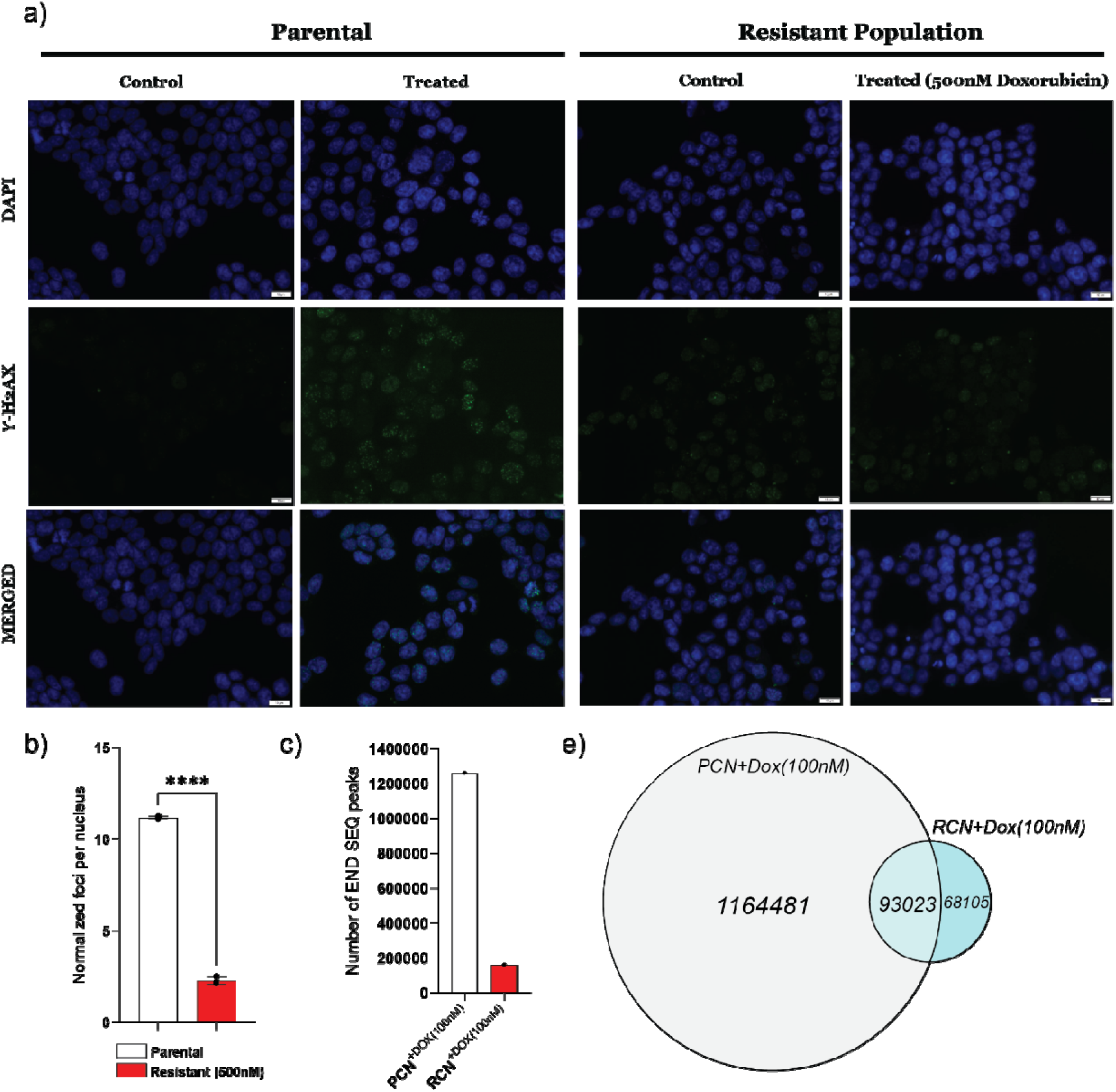
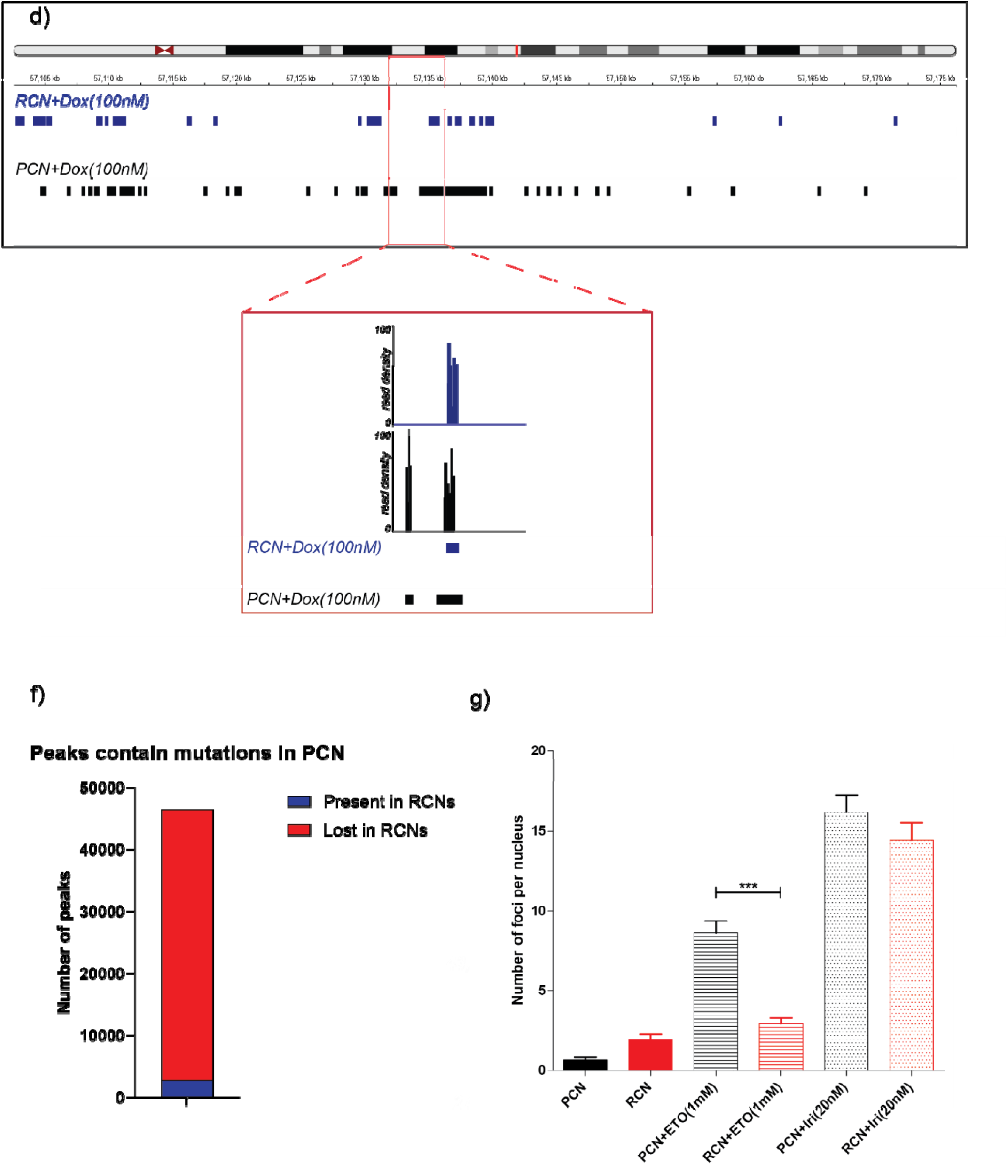
The adapted population experiences a lower frequency of DSB following Dox exposure. (a) DSBs were visualized by the formation of γH2AX foci in cells exposed to Dox (100 nM for 6 h). PCN cells were compared to the RCN1 clone adapted to 500 nM Dox. Experiments were conducted in triplicate. (b) Quantification of data presented in (a) showing the number of foci generated upon Dox-treatment in PCN and RCN1 clones (n = 368). Images were analyzed with the integrated software image are at a scale of 12.5 µm. The significance of differences was determined using an unpaired Student t-test (**** p < 0.00021) denoted above in (b). (c) Bar graph showing the number of DSB sites in doxorubicin-sensitive (PCN^+Dox(100nM)^) and doxorubicin-resistant (RCN1^+Dox(100nM)^) cell populations following treatment with 100□nM doxorubicin for six hours. (d) A representative example of the distribution of the END-seq peaks in PCN+Dox^100nM^ (black) and RCN+Dox^100nM^ (blue) along chromosome 14. The red highlighted region provides a zoomed-in view of specific loci on chromosome 14, illustrating changes in read density at the break sites. Read density in RCN cells treated with 100 nM Dox is shown in blue (top), while PCN cells under the same treatment condition are shown in black (bottom). (e) A Venn diagram illustrates the total number of END-seq peaks identified in PCN+Dox^100nM^ and RCN+Dox^100nM^ samples, as well as the number of shared peaks between the two groups. (f) The stacked bar graph highlights the number of END-seq peaks containing mutations in PCN (see in detail figure 2b legend). A large proportion of these peaks were lost in RCN (red), while the remaining peaks (blue) were retained in both PCN and RCN. (g) Doxorubicin resistance reduces the ability of etoposide, but not SN-38, to generate DSB. PCN and RCN cells were treated with Etoposide (1μM, 1h) and SN-38 (20nM, 1h), and γH2AX foci were assessed by immunofluorescence.

To distinguish between a global reduction in Dox activity (e.g., decreased intracellular drug levels) and a site-specific loss of Top2-mediated cleavage, we performed END-seq to map genome-wide DSBs in PCN and RCN1 cells. Consistent with the γH2AX data, resistant cells showed an approximately 10-fold reduction in the number of Dox-induced DSB sites compared to parental cells (Fig. 5c). Importantly, the remaining DSB peaks in RCN1 cells largely overlapped with or were located near peaks observed in PCN cells (Fig. 5d,e), suggesting that these sites either remained unmutated or retained the ability to support Top2-mediated cleavage.

Crucially, many genomic regions that exhibited DSB peaks in PCN cells but not in RCN1 cells corresponded to sites that had acquired mutations during adaptation (Figs. 2b–d, 5f). This strong spatial association indicates that mutation of Top2 cleavage sites directly contributes to the loss of DSB formation upon subsequent Dox exposure, thereby promoting resistance.

The model further predicts that resistance driven by the loss of Top2-mediated cleavage should extend to other Top2 inhibitors but not to agents that induce DNA damage through distinct mechanisms. To test this, we compared γH2AX formation in response to etoposide (another Top2 inhibitor) and SN-38 (the active metabolite of the Top1 inhibitor irinotecan). Consistent with our hypothesis, RCN cells displayed reduced γH2AX foci following etoposide treatment compared to PCN cells, whereas no reduction was observed following SN-38 treatment (Fig. 5g, Fig. S9a,b).

Together, these results provide strong experimental support for the proposed mechanism: accumulation of mutations at Top2 cleavage sites reduces the ability of Dox to induce DNA damage, thereby conferring resistance while preserving sensitivity to agents that act through distinct pathways.

### Cells resistant to Dox lose the ability to adapt to Top1 inhibition

To our surprise, unlike DSB induced upon Dox treatment, the γH2AX assay demonstrated a significantly higher number of DSBs in untreated resistant clones compared to the untreated parental clone (Fig. 6a). We suggested that such an increase in the baseline DSB levels in the resistant cells could result from impaired relaxation of topological stress during transcription or replication resulting from the reduction in the number of functional Top2 binding sites during Dox-induced selection. Accordingly, analysis of breakage signals in resistant and sensitive cells across genes revealed that RCN cells exhibited increased breakage in gene bodies (Fig. 6b). Furthermore, breakage levels correlated with gene expression levels, whereas highly expressed genes showed higher levels of breakage compared to genes with low expression levels (Fig. 6b,c). Since gene bodies are hotspots for topological stress, these findings suggest that the increased fragility in untreated RCN cells arises from excessive topological stress and potentially a greater reliance on Top1.

**Figure 6.**
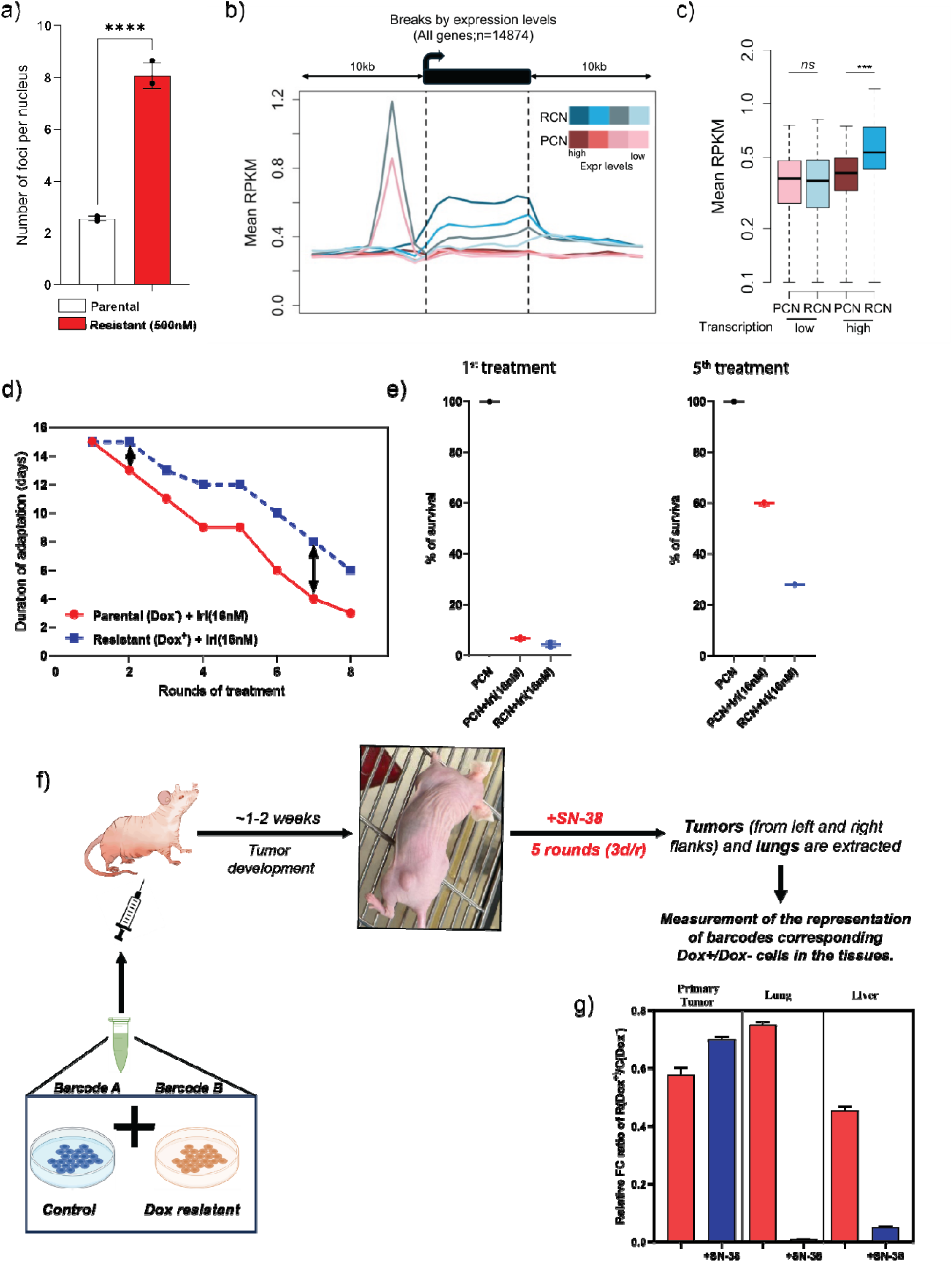
Impact of Dox resistance on the acquisition of resistance to irinotecan. (a) Quantification of gH2AX foci in naïve resistant clones compared to naïve parental cells. Statistical significance was assessed using an unpaired Student’s *t*-test (**** *p* < 0.0001), as indicated in panel (a). (b) END-seq signal across gene bodies and 10kb flanking regions in RCN (blue shades) and PCN (red shades). Lines represent the mean RPKM (Reads Per kilobase of transcript per Million mapped reads) at each gene’s quartile grouped by expression levels. (c) Boxplots representing the average RPKM over gene body in low (first quartile) and high (fourth quartile) groups of genes in resistant and sensitive cells. T-test ***p<1e-10 (d) The adaptation to irinotecan timelines of control and Dox-resistant cells. A total of 8 cycles of treatment with 16nM irinotecan of PCN and RCN cell populations were applied (e) The Boxplot showing the adaptation of PCN and RCN1 to 16nM irinotecan. PCN and RCN1 cells underwent five treatments with irinotecan or remained untreated. Cell survival in these cell populations following treatment with 16nM irinotecan was measured by XTT. (f) Graphical demonstration of *in vivo* experiment designed for testing the differences in sensitivity to irinotecan of PCN and RCN1 cells. (g) The results of the mouse experiment are illustrated in Fig. 5f. The data shows the ratio of representation of barcodes corresponding to PCN and RCN1 in the primary tumors and metastasis in lungs and livers. Treatment was done twice a week in 5 cycles

Based on this, we reasoned that inefficient DNA relaxation by Top2 may render Dox-resistant cells more dependent on DNA relaxation by Top1. In such a scenario, Top1 inhibition may impose greater toxicity on these cells, potentially preventing them from developing resistance to Top1 inhibitors, since resistance to Top1 inhibitors is similarly associated with a gradual loss of Top1-binding sites on DNA, and therefore a gradual loss of Top1-dependent DNA relaxation [18]. To explore this possibility, both parental PCN and Dox-resistant RCN1 cells were exposed to eight cycles of 16nM of a Top1 inhibitor SN-38 (an activated component of the pro-drug Irinotecan) for 48 hours and the recovery time was measured following each cycle. Though the original response of PCN and RCN cells to SN-38 was similar, we found that Dox-resistant clones adapted to the Top1 inhibitor significantly more slowly than the parental clone (Fig. 6d). Specifically, while PCN and RCN1 cells showed similar sensitivity to SN-38 in the first treatment cycle (around 6% survival), survival of PCN cells increased to 60% by the fifth treatment cycle, while survival of RCN1 cells increased to about 25% (Fig. 6e). Thus, the development of Dox resistance led to slower adaptation and, eventually, higher sensitivity to irinotecan.

Next, we investigated whether the primary tumors and metastasis generated by Dox-resistant RCN1 cells in nude mice exhibited increased sensitivity to irinotecan therapy compared to the parental PCN cells. For these in vivo experiments, we used cell barcoding as described earlier [38]. PCN and RCN1 cell populations were barcoded differently, mixed, and then injected into mice subcutaneously. Following one week of tumor formation, the mouse groups underwent treatment with irinotecan over five sessions, with each administration occurring every three days (Fig. 6f). Upon completion of the fifth treatment, primary tumors, livers, and lungs were collected. DNA was isolated from these tissues, and the barcodes were quantified by RT-PCR. The subsequent quantitative analysis compared the ratio of PCN and RCN1 cells in both primary tumors and the metastases in the lungs and liver. While the ratio of PCN and RCN1 cells did not change upon irinotecan treatments in the primary tumor, the RCN1 cells were significantly underrepresented in both lung and liver metastasis in irinotecan-treated animals (Fig. 6g), indicating that they were significantly more sensitive to irinotecan.

In summary, outcomes from experiments conducted both *in vitro* and *in vivo* suggest that cancer cells adapting to multiple exposures to Dox showed reduced adaptation to exposures to the Top1 inhibitor irinotecan, and accordingly increased sensitivity to this drug, which could have significant implications in clinical settings, leading to novel treatment strategies for Dox-resistant patients.

## DISCUSSION

In this study, we demonstrate a novel mechanism of Dox resistance and provide a model to explain the need of multiple exposures to drugs and dose escalation for drug-resistance selection. Such a selection strategy implies that cells may have adapted to low drug concentrations, potentially through mechanisms such as epigenetic changes, guiding further selection of resistant mutant forms. Alternatively, the development of drug resistance may arise through progressive accumulation of a large number of mutations, each providing a fraction of the resistance, which are subsequently selected during the process of dose escalation. Our study supports the latter model by demonstrating that a very high number of mutations were generated in the process of the selection of Dox-resistant mutants in the dose escalation *in vitro* experiment.

Specifically, we observed that sequential treatments of cancer cell lines with escalating doses of Dox led to progressive accumulation of very specific mutations. Mathematical analysis presented in Materials and Methods and Supplementary Materials demonstrates that the probability of random generation of these mutations is nearly zero, indicating that they are not random and must be guided by a special mechanism. There are several lines of evidence that link these mutations to Top2-induced cleavage sites: (a) the majority of these mutations result from the repair of DSBs via HR and NHEJ systems, (b) the mutations co-localize with Dox-induced DSBs, as measured by END-seq, (c) in Dox-resistant cells the majority of Dox-induced DSBs that overlap with the mutations are lost, (d) a significant fraction of the CTCF binding sites overlap with the mutations.

We argue that accumulation of the Dox-induced mutations hinders Top2 activity at these sites during subsequent exposures to the drug, ultimately leading to the development of Dox resistance. It should be emphasized that Top2 generates DSBs specifically at the location of Dox DNA intercalation, which is strongly dependent on the chromatin architecture and is linked to epigenetics and chromatin opening [36]. Therefore, to reduce the generation of DSBs and achieve resistance, it is not necessary to inactivate the majority of Top2 cleavage sites, but only sites adjacent to the intercalated dox molecules.

An important question that could be raised is how these findings related to numerous prior studies on the Dox resistance, which identified a set of mutations and changes in the transcriptome associated with the resistance [21,27,28,30,32]. In our system, we did not find any of these previously reported mechanisms. Specifically, we found no evidence of MDR pump overexpression, Top2 downregulation or mutations, or decreased expression of apoptotic proteins (Table S1, S2). It is possible that these alterations emerge at late stage of Dox adaptation or represent parallel resistance mechanisms. For example, part of Dox effects is related not to Top2-mediated DNA cleavage, but to the eviction of histones from nucleosomes [36] These effects could lead to significant changes in the transcriptome, including changing the balance towards better Dox resistance.

Surprisingly, we found that while the number of Dox-mediated DSBs was significantly lower in the resistant clones compared to the parental clone, spontaneous DSBs accumulated at higher levels in the resistant cells. These data suggested that DSB in untreated cells may be caused not by the function of Top2 or Top1, but rather by unresolved topological stress during replication or transcription. Such stress could be caused by the insufficiency of Top2 activity due to the mutations in a large fraction of Top2 binding sites.

Based on these findings, we suggest that the resistant cells may be more dependent on the Top1 activity for DNA relaxation during transcription compared to normal cells. Accordingly, RCN cells may become more sensitive to inhibitors of Top1 and may constrain the ability of cells to adapt to Top1 inhibitors such as irinotecan. Recently, we found that adaptation to irinotecan involves the generation of multiple mutations in the Top1 binding sites [18,19]. Therefore, Dox-resistant cells that lost a large fraction of Top2 binding sites may not be able to develop resistance to irinotecan since they may not survive reduced Top1 activity. In line with that, monitoring the dynamics of adaptation to irinotecan in PCN and RCN clones revealed that such an adaptation was significantly delayed in the RCN cells, and they retained irinotecan sensitivity significantly longer.

Furthermore, an *in vivo* experiment with multiple exposures to irinotecan uncovered that metastasis derived from the RCN cells were significantly more sensitive to irinotecan than metastasis derived from PCN cells. Overall, our experiments conducted both *in vitro* and *in vivo* revealed that cells acquiring resistance to Dox concurrently develop sensitivity to irinotecan. This finding provides a mechanistic explanation for the significant efficacy of irinotecan and topotecan following the doxorubicin failure in relapsed Willms tumors and Small Cell Lung Cancer [39]. Similarly, irinotecan has been efficiently used as salvage therapy following failure of doxorubicin in Neuroblastoma [40]. This suggests that irinotecan could serve as a compelling salvage therapy for other cancers, as well.

## 4. MATERIALS AND METHODS

### 4.1 Cell culture and reagents

HCT116 cells (ATCC Cat# CCL-247, RRID:CVCL_0291) were obtained from the American Type Culture Collection and maintained in McCoy’s 5A medium supplemented with 10% fetal bovine serum (FBS), 4 mM L-glutamine (BI-Biologicals, Cat#03-020-1B), 2 mM L-alanyl-L-glutamine (BI-Biologicals, Cat#03-022-1B), and 1% penicillin-streptomycin (BI-Biologicals, Cat#03-031-1B). H1975 cells (NCI-H1975) were obtained from the American Type Collection and maintained in RPMI-1640 medium supplemented with 10% FBS, 4 mM L-glutamine, 2 mM L-alanyl-L-glutamine, and 1% penicillin-streptomycin. Cells were incubated at 37□°C in a humidified atmosphere containing 5% CO□. Doxorubicin and SN-38 were sourced from Sigma-Aldrich (St. Louis, MO, USA).

### 4.2 Cell cloning

To establish monoclonal populations, cells were subjected to limiting dilution cloning. Individual clones were selected using cloning discs (Merck, Darmstadt, Germany, Cat# Z374431), then expanded and cryopreserved in liquid nitrogen in a freezing solution composed of 10% dimethyl sulfoxide (DMSO) in fetal bovine serum (FBS) for long-term storage.

### 4.3 Cell Viability assay

Cytotoxicity of doxorubicin was determined using the XTT assay [41]. HCT116 and H1975 cells were seeded in 96-well plates and allowed to recover. The next day, cells were treated with the drug at concentrations ranging from 25 to 800 μg/mL, in triplicate, for 24 h incubation. 50 μL XTT (5 mg/mL) was added to each well. After 2 h of incubation, absorbance was determined using a spectrophotometer at 450 nm. To measure non-specific signal, a 630–690 nm wavelength was used.

### 4.4 Crystal Violet assay

The Crystal Violet (CV) assay [42] was performed according to the protocol outlined by different studies. The staining solution consists of 0.125g crystal violet powder and 20% ethanol. 200 mL Lysis solution (0.1M sodium citrate, 5.88g, 50% ethanol 100 mL, dH2O 100 mL, pH 4.2 adjusted with HCl) was used. Cells were seeded in a 6-well plate, washed with PBS. Followed by the addition of 150 µL of staining solution to each well. Incubation was performed for 30 minutes at room temperature. 350 µL lysing solution was added and incubated for 30 min. 60 µL of the sample was diluted 5 times in DDH2O, followed by measurement of OD at 590 nM.

### 4.5 DNA barcoding

Barcoding of the cloned cell population was performed using the Cellecta CloneTracker 50M Lentiviral Barcode Library (RRID:SCR_021827), in accordance with the manufacturer’s protocol. For animal experiments, cells, home-made DNA barcoding libraries were used [38]. Cells were transduced with the lentiviral barcoding library and subjected to puromycin selection to enrich for successfully infected cells. Following selection, cells were allocated into separate groups for drug treatment. After recovery, genomic DNA was extracted, and the integrated barcodes were amplified via nested PCR. All resulting samples were pooled and prepared for high-throughput sequencing.

### 4.6 DNA extraction and amplification of library barcodes, qPCR

Genomic DNA was extracted from cultured cells using the Wizard Genomic DNA Purification Kit (Promega, Madison, WI, USA). Barcode sequences were amplified via nested PCR. The first PCR (PCR1) utilized Titanium Taq DNA Polymerase (Takara Bio, Cat# 639209), and PCR products were purified using the QIAquick PCR & Gel Cleanup Kit (Qiagen, Hilden, Germany). A second round of amplification (PCR2) was performed with nested primers—either generic or sample-specific—using the Phusion High-Fidelity PCR Master Mix (Thermo Scientific, Waltham, MA, USA). A secondary barcode was added during PCR2 to enable sample multiplexing. PCR products were normalized individually, pooled, and further purified using AmpureXP magnetic beads (Beckman Coulter, Brea, CA, USA) according to the manufacturer’s instructions. Barcode abundance specific to PCN (parental) and RCN (resistant) cells was quantified using qPCR.

### 4.7 RNA sequencing and transcriptome analysis

Total RNA was extracted using the RNeasy Mini Kit (Qiagen, Cat#74104). Library preparation and sequencing were performed by BGI (China) using the BGISEQ-500 platform (RRID:SCR_017979). Raw sequencing reads were processed to remove adapter sequences and low-quality reads. Read quality was assessed using FASTQC [43] based on Phred quality scores. High-quality reads were aligned to the human reference genome (hg38) using HISAT2 [44] with default settings. Reads were then imported into SAMtools [45] for conversion into BAM files. Gene-level quantification was conducted with featureCounts [46]. To enable an accurate comparison of gene expression between parental and drug-treated clones, count data were normalized using the TMM (trimmed mean of M-values) method implemented in the edgeR R package [47]. For differential expression analysis, data were transformed using the voom method, and linear modeling was performed with the limma package [48]. Multiple testing correction was applied using the Benjamini–Hochberg (BH) procedure, and genes with a false discovery rate (FDR) < 0.05 and a fold change > 1 were classified as differentially expressed. Genes were ranked based on fold change values and analyzed using Gene Set Enrichment Analysis (GSEA) [49] to identify significant signaling pathways altered during drug treatment. Enrichment analysis was performed against the hallmark gene sets from the Molecular Signatures Database (MSigDB), version 7.2 [50].

### 4.8 Whole-Genome Sequencing (WGS) analysis

Genomic DNA samples were submitted to Dante Labs (Italy) for whole-genome sequencing at 30× coverage. Upon receipt of sequencing data, quality control of raw reads was performed using FastQC to assess base quality, adapter content, and overall sequence integrity. Demultiplexing of reads was conducted using barcode-splitter (V.0.18.6), allowing a single mismatch or deletion within barcode sequences. High-quality reads were aligned to the human reference genome (build hg38) using BWA-MEM [51] with default parameters. Duplicate reads were removed using Picard. The aligned SAM files were converted to BAM format using samtools.

Variant calling was performed using the Genome Analysis Toolkit (GATK) in accordance with the GATK Best Practices workflow [52], generating combined variant call format (VCF) files. Local realignment and base quality recalibration were performed using default parameters. Subsequent data visualization and statistical analysis were performed in R using the dplyr (47) and karyoploteR (48) packages. Functional annotation of variants was conducted using SnpEff [53]. To match detected variants to known genomic features, bedtools was used in conjunction with BED files from UCSC, ENCODE databases [54,55]. Metadata extraction and comparative evaluation were performed using bcftools in the Linux environment. Raw data and corresponding VCF files are publicly available through the NCBI Sequence Read Archive (SRA) under accession number.

#### 4.8.1 The Mutations Resonance Analysis

To assess whether the observed overlap of mutations among independently derived clones could arise by chance, we performed a combinatorial probability analysis (see in detail Supplementary file “The Mutation Resonance Problem”).

m = 11 × 10^3^ – denote the total number of mutations detected in each clone in one cycle of treatment.

n = 3 × 10^9^ – total size of the human genome in bp.

k – shows the number of shared mutations across clones.

##### For mutations present in two out of three clones

k = ∼1680 – mutations were shared in two clones in one treatment cycle.

The probability of two randomly chosen subsets of size *m* from a universal set of size *n* having an intersection of size *k* is given by:

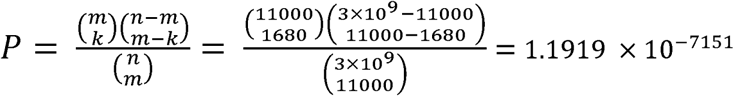

##### For mutations present in all three clones

k = 7600 – is the number of mutations shared across all three clones in one treatment cycle.

The probability of three randomly chosen subsets of size *m* from a universal set of size *n* having an intersection of size *k* is.

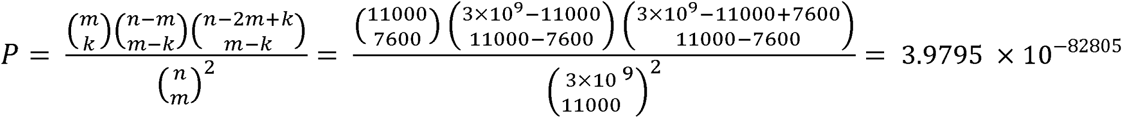

These astronomically small probabilities indicate that the observed overlaps are highly unlikely to occur by random chance. Therefore, the data strongly reject the null hypothesis of independent mutation generation and suggest the presence of either strong selection or directed mechanisms that shape the mutational landscape in these clones.

### 4.9 DNA breaks quantification and identification, γH2AX Assay

Parental PCN cells and Doxorubicin-adapted cell lines (RCN1, RCN2, RCN3) were treated with 100 nM Doxorubicin for 6 hours. Cells were washed with 1× PBS and fixed using 1% formaldehyde for 5 min at room temperature, followed by permeabilization with 0.2% Triton X-100 in PBS for 10 minutes at room temperature. Blocking was performed with 5% (w/v) bovine serum albumin (BSA) in PBS containing 0.05% Tween-20 (PBST) for 1 hour. Cells were washed with PBST 3 times and incubated overnight at 4□°C with a primary antibody against phospho-Histone H2A.X in 3% BSA (Ser139) (Cell Signaling Technology, Cat# 2577, RRID: AB_2118010). After five PBST washes, a secondary antibody (Goat Anti-Rabbit IgG H&L, Alexa Fluor 488; Abcam Cat# ab150077, RRID: AB_2630356) was applied for 1 hour. Non-specific binding was removed by additional PBST washes. Imaging was performed using the Hermes WiScan High Content Imaging System (IDEA Bio-Medical, RRID: SCR_021786), and image analysis was carried out with Athena Wisoft software (Ver1.0.10), using fixed maxima (550) for foci detection and automatic background correction based on untreated control. Statistical analyses were conducted using the one-way ANOVA statistical method. The significance of the differences was determined using an unpaired two-tailed t-test.

### 4.10 END-seq

Human colon cancer cells (HCT116) were cultured in McCoy’s 5A medium supplemented with 10% fetal bovine serum (FBS, Gibco), penicillin-streptomycin, L-Glutamine, and Sodium Pyruvate (Gibco) at 37 °C in a 5% CO2 atmosphere. HCT116 cells were treated with doxorubicin (85nM) for 3h. For END-seq treatment, cells were synchronized at G1 phase using a 22-hour treatment with the CDK4/6 inhibitor Palbociclib (5 µM). To validate cell cycle synchronization, cells were fixed in 100% ethanol, treated with RNase A (1 mg/mL), and stained with propidium iodide (50 µg/mL). Cell cycle distribution was analyzed using a Gallios Flow Cytometer. END-seq has been conducted as previously described [56]. Briefly, single-cell suspensions of synchronized parental and DOX-resistant HCT116 cells (7 million), were washed in PBS, and resuspended in cell suspension buffer (Bio-Rad), embedded in low-melting agarose, and transferred into plug molds (Bio-Rad CHEF Mammalian Genomic DNA plug kit). Plugs were allowed to solidify at 4°C and were then incubated with Proteinase K solution (Puregene, QIAGEN) for 1h at 50°C and then for ON at 37°C, followed by consecutive washes in a wash buffer containing 10 mM Tris-HCl (pH 8.0) and 50 mM EDTA (Wash Buffer) and then in a TE Buffer containing (10 mM Tris (pH 8.0) and 1 mM EDTA). Washed plugs were subsequently treated with RNaseA (Puregene, QIAGEN), washed again in Wash Buffer, and stored at 4°C (up to 2-4 days). Blunting, A-tailing, and ligation to biotinylated hairpin adaptor 1 were done at 37°C, 4°C, respectively. DNA recovered from melted plugs was sheared to a length between 150 and 200 bp by sonication (Covaris), and biotinylated DNA fragments were purified using streptavidin beads (MyOne C1, Invitrogen). Following streptavidin capture, the newly generated ends were end repaired using T4 DNA polymerase (15 U), Klenow fragment (5 U), and T4 polynucleotide kinase (15 U); A-tailed with Klenow exo-fragment (15 U); and finally ligated to hairpin adaptor 2 using the NEB Quick ligation kit. After the second adaptor ligation, libraries were prepared by first digesting the hairpins on both adapters with the USER enzyme (NEB) and PCR amplified for 18 cycles using TruSeq index adapters. All libraries were quantified using qPCR. Sequencing was performed either on the Illumina Nextseq550 (75 bp single-end reads).

#### 4.10.1 END-seq downstream analysis

Raw FASTQ files were first assessed for their quality using FastQC [57], followed by adapter trimming using the Perl wrapper Trim Galore tool [58]. High-quality reads were then aligned to the human reference genome (hg38) using Bowtie2 [59]. The resulting SAM files were converted to BAM format and sorted using Samtools [45]. To ensure accurate downstream analysis, Picard’s MarkDuplicates tool was employed to mark PCR duplicates, as well as low-quality and multimapped reads. MACS2 [60] was then used for peak calling to identify genomic regions enriched for DNA double-strand breaks (DSB). Only peaks supported by at least 40 reads were retained for further analysis. Finally, correlation analysis was performed between the identified total number of DSB peaks (1.2 million) and common mutations observed across single-cell clones.

### 4.11 In vivo experiment

All animals were housed under pathogen-free conditions, and the Ariel University Animal Care and Use Committee approved all animal procedures. Cell line suspensions were prepared in 1:1 Matrigel (ECM Gel #E1270, Sigma-Aldrich), and approximately 1.5 × 10^6^ cells were subcutaneously injected into both flanks of 9-week-old female nude (Nu/Nu) mice (Envigo, Ness Ziona, Israel). Tumors were measured with calipers, and the tumor volume was calculated according to the formula Vol = 0.52 × L × W2. Two groups, control and treated (with 2 mice per group), were assigned for the experiment. When tumors reached the volume of approximately 200-300 mm³, control groups were euthanized, whereas treated groups were assigned to treatment with Irinotecan, a Top1 inhibitor. Irinotecan was administered intraperitoneally twice a week for 5 rounds. For treated mice, tumor volumes were measured, and body weights were monitored twice weekly for the entire span of the experiment. After the experiment was finalized, mice were sacrificed, and primary tumors and metastasized organs (lung, liver) were collected.

### 4.12 Statistical Analysis

Statistical analysis was performed either in the R programming language or GraphPad Prism. Data are expressed as mean ±SD. Student’s t-test and one or two-way analysis of variance (ANOVA) were used to evaluate the statistical significance of the difference in two or more groups, respectively. An empirical P-value was calculated based on a non-parametric Monte Carlo permutation test. A P-value of less than 0.05 was considered significant. ns, not significant; ∗p <0.05, ∗∗p < 0.01, and ∗∗∗p < 0.001, ∗∗∗∗p < 0.0001.

## Data Availability Statement

All raw sequencing data generated in this work are stored under BioProject **PRJNA1507608**. The processed raw sequencing data were provided in the main manuscript and supplementary files.

## Acknowledgments

This work was supported by grants from the United States–Israel Binational Science Foundation (BSF; Grant No. 2021137 to Michael Y. Sherman), the Israel Science Foundation (ISF; Grant No. 731/24 to Michael Y. Sherman and Grant No. 1920/20 to Yaakov Maman), the Ministry of Education, Israel (Grant No. 1001828974 to Michael Y. Sherman), and the Israel Cancer Research Fund (ICRF; Grant No. 23–201-RCDA to Yaakov Maman).

