## Supplementary Figures and legends, Tables for "Systematic Generation of Mutations at Topoisomerase II Cleavage Sites Enables Cancer Adaptation to Doxorubicin Therapy": SUPPLEMENTAL FIGURES and LEGENDS.docx

SUPPLEMENTAL FIGURE LEGENDS

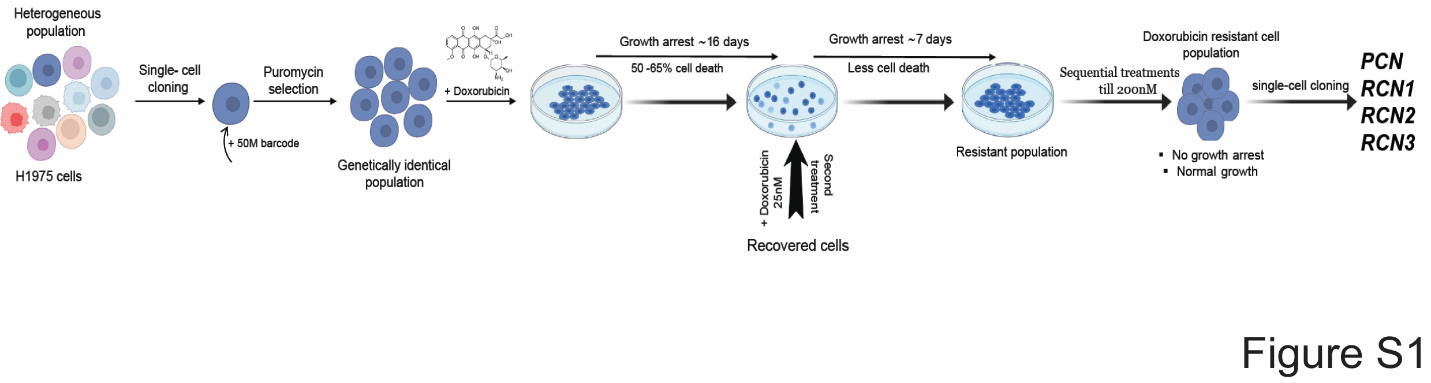

**Supplementary Figure S1**. A graphical representation of the experimental plan, showing dose selection of Dox-resistant H1975 cells (RCN1, RCN2, RCN3).

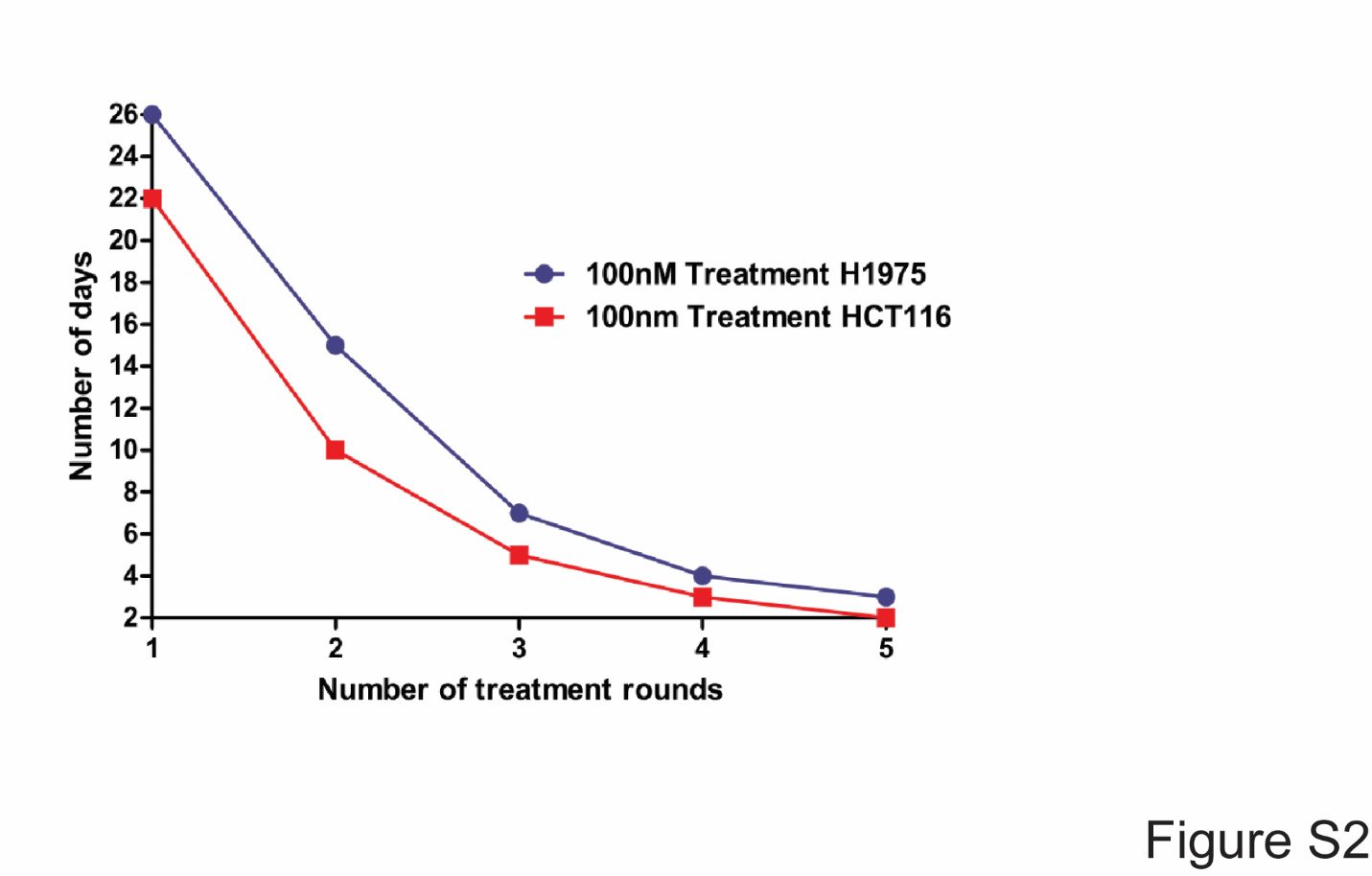

**Supplementary Figure S2**. The line plot illustrates the response kinetics of H1975 and HCT116 cells to repeated treatments with 100 nM doxorubicin over five consecutive rounds. With each treatment cycle, both cell lines exhibited progressively increased resistance, demonstrating a faster recovery and reduced sensitivity to the drug.

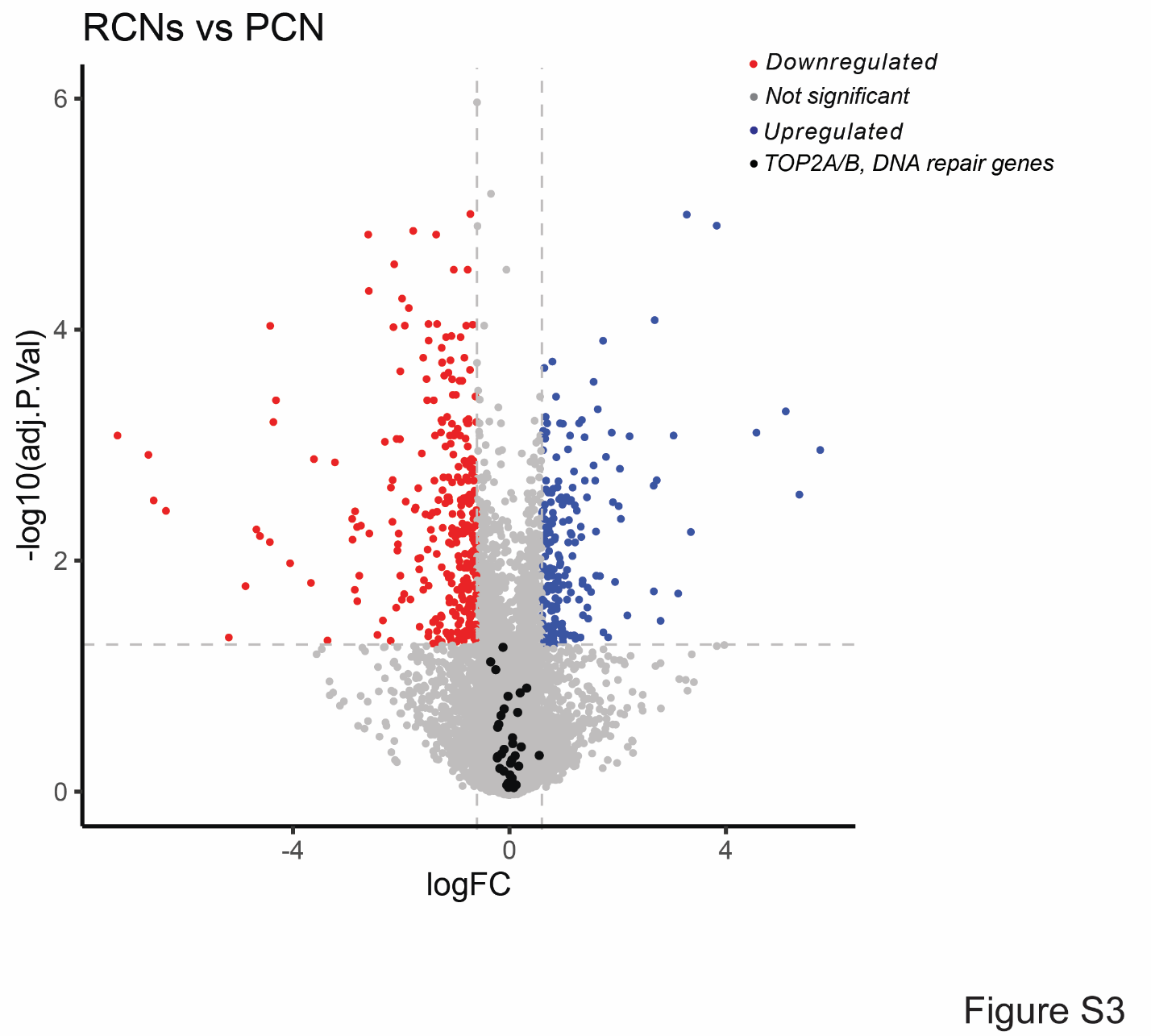
**Supplementary Figure S3.** The Volcano plot shows the differentially expressed genes in comparison of RCN (resistant clone) vs PCN (parental clone).

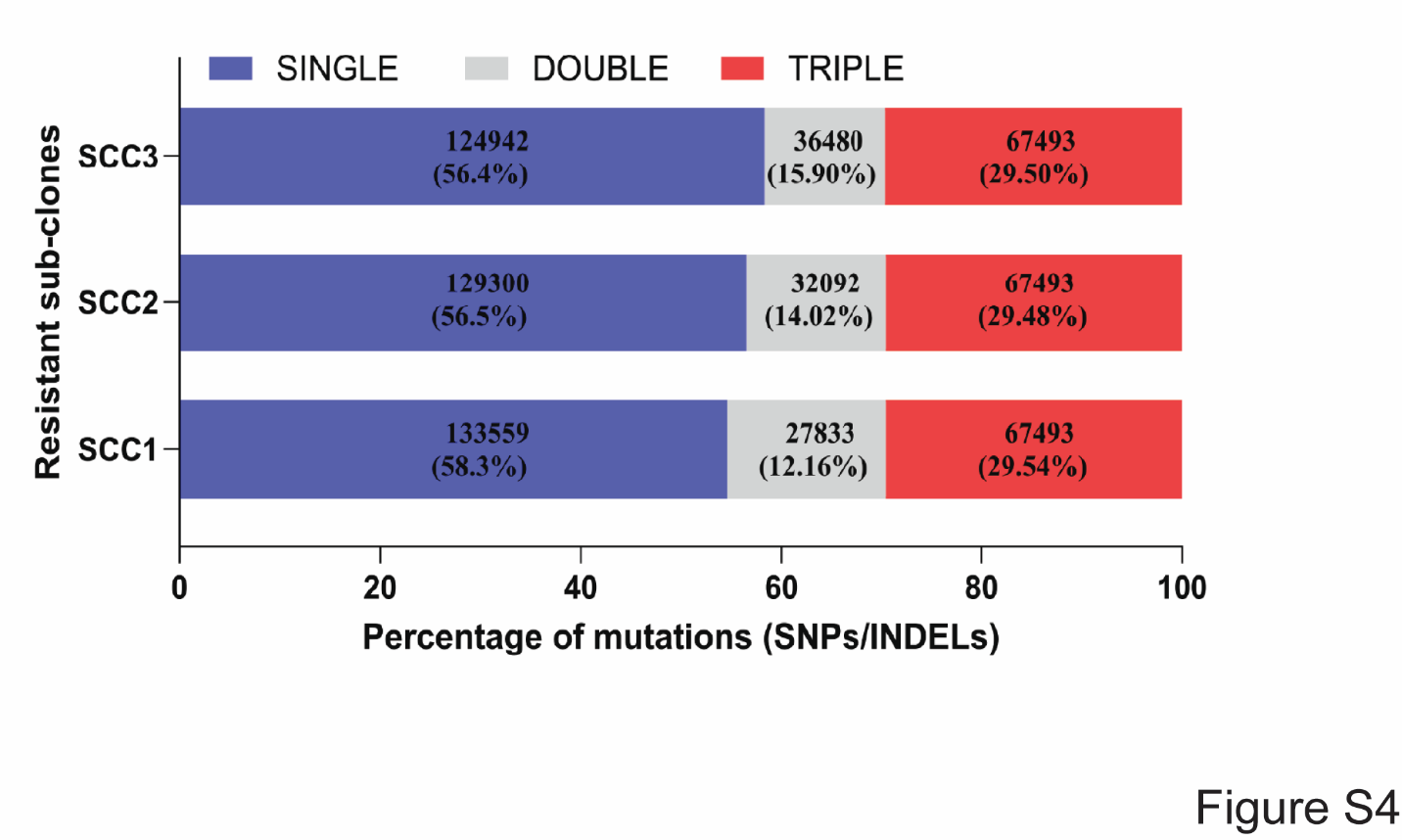

**Supplementary Figure S4.** Quantification of overlapping mutations in doxorubicin-resistant H1975 clones emerging during the adaptation process. The stacked bar graph displays the distribution of shared and unique mutations among the three resistant clones. Mutations are categorized as follows: blue indicates clone-specific mutations that are not present in the other clones; red represents mutations shared by two out of three clones; and green denotes mutations common to all three clones.

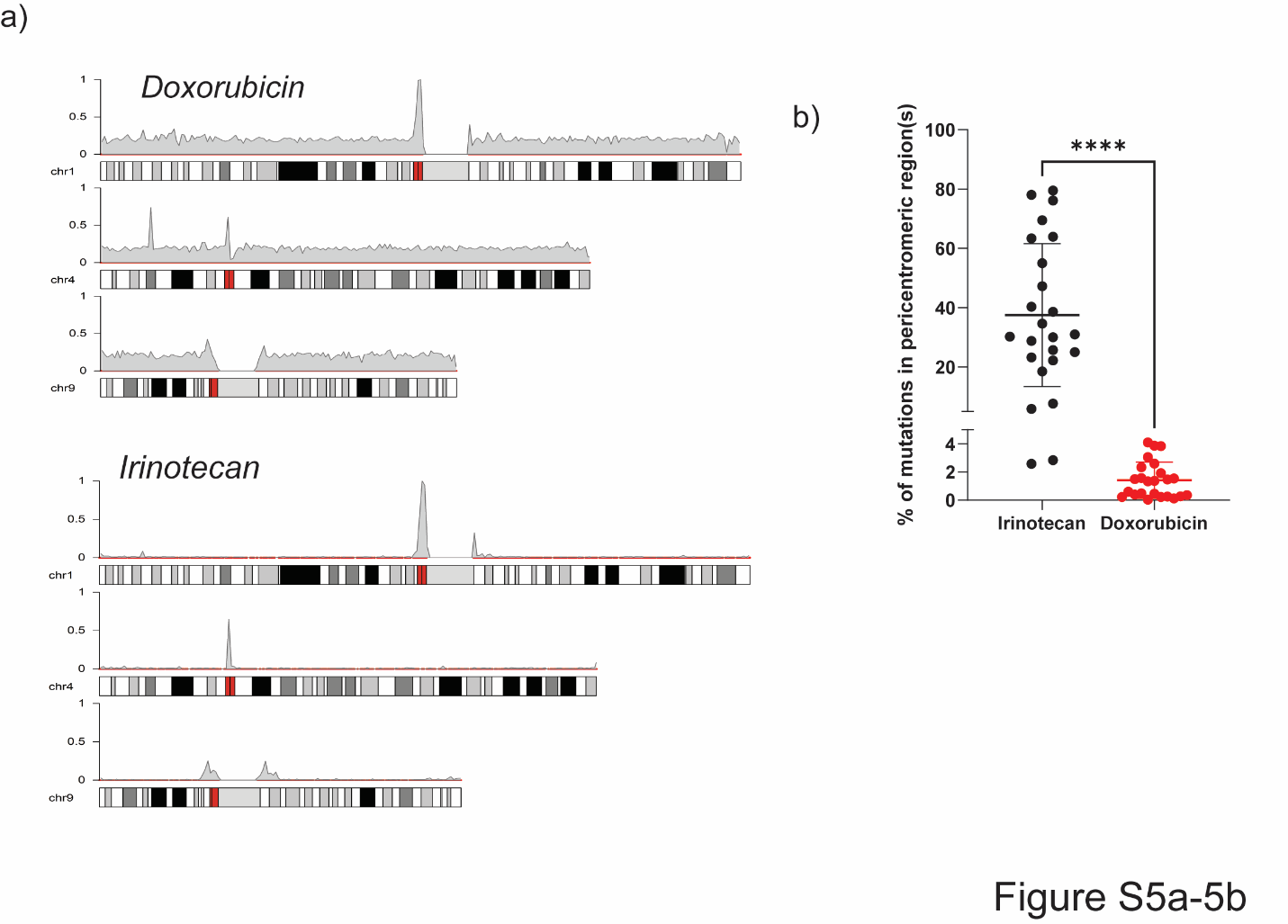

**Supplementary Figure S5.** (a) Comparison of the distribution of overlapping mutations in Dox-resistant and irinotecan-resistant cells (Kumar et al., 2023) along chromosomes 1, 4, and 9, shown as representative examples. (b) A scatter plot illustrates differences in the accumulation of mutations within the pericentromeric regions across chromosomes in Irinotecan and Doxorubicin-treated samples. Each dot represents a single chromosome (22 autosomes plus X and Y), with the Y-axis indicating the percentage of mutations localized to the pericentromeric region for that chromosome. The significance of differences was determined using an unpaired Student t-test (**** p < 0.0001)

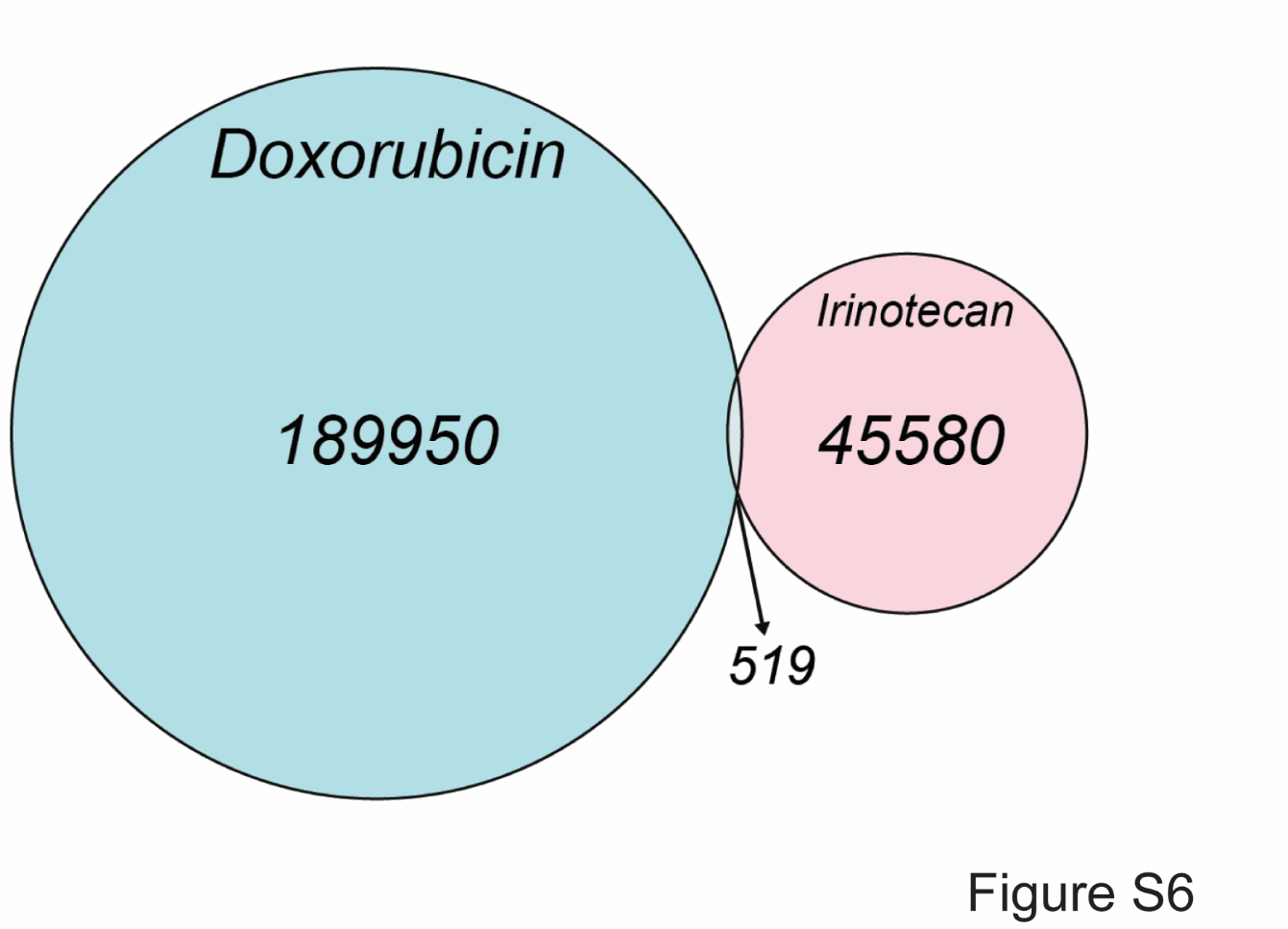

**Supplementary Figure S6.** A Venn diagram illustrates the overlap of mutations identified in HCT116 clones following treatment with doxorubicin and irinotecan, respectively. The limited number of shared mutations highlights the distinct mutational profiles induced by each drug.

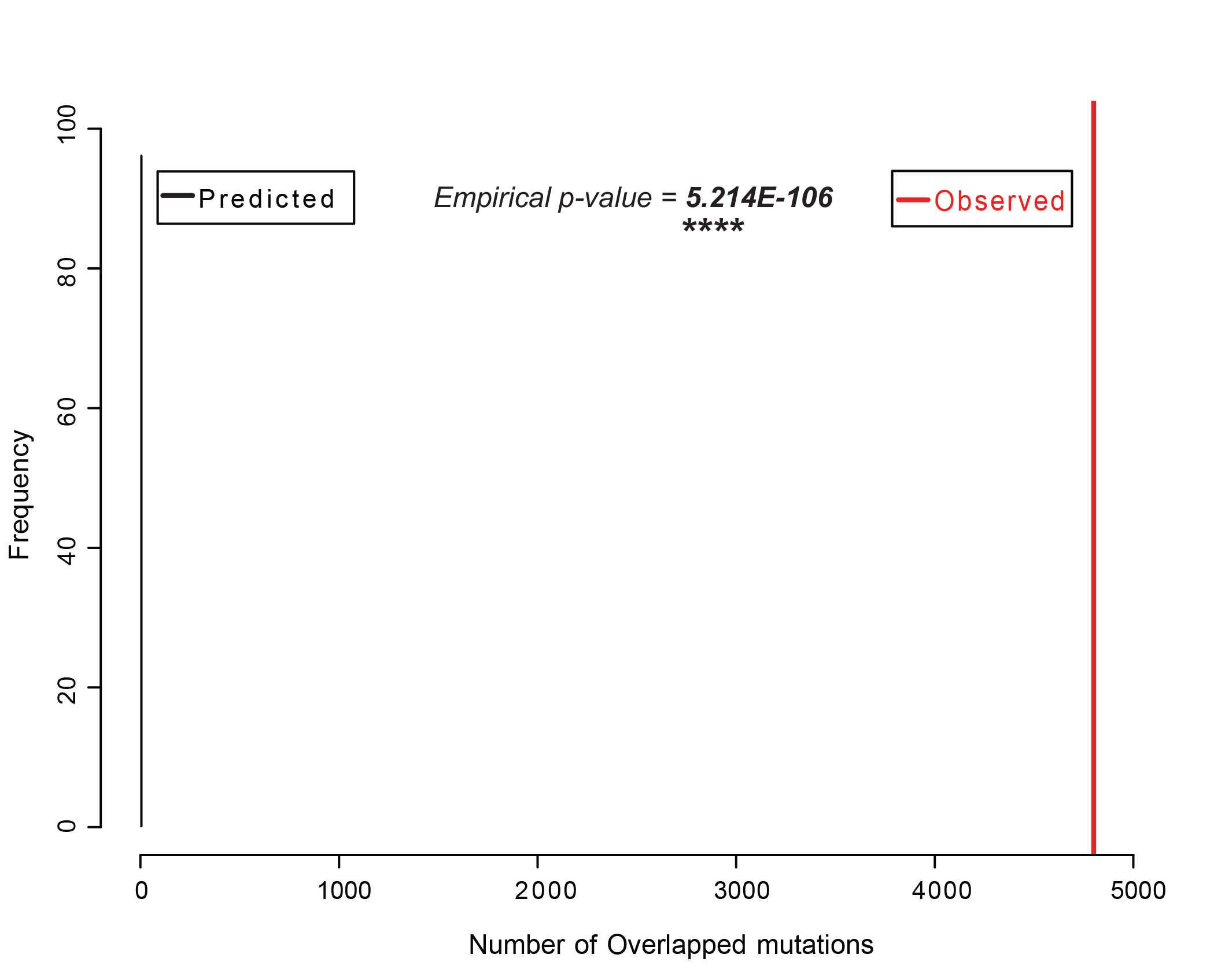

**Supplementary Figure S7.** The permutation plot illustrates the statistical comparison between the observed and expected overlap of common mutation sites with Top2-associated binding regions. The black line represents the overlap values obtained from 100 permutations of randomly generated Top2 binding sites across the genome. The red line indicates the actual observed overlap with the mutation dataset. A Monte Carlo permutation test (100 iterations) was used to assess statistical significance. The resulting p-value reflects whether the observed overlap is greater than expected by chance. The y-axis indicates the number of permutations contributing to the null distribution.

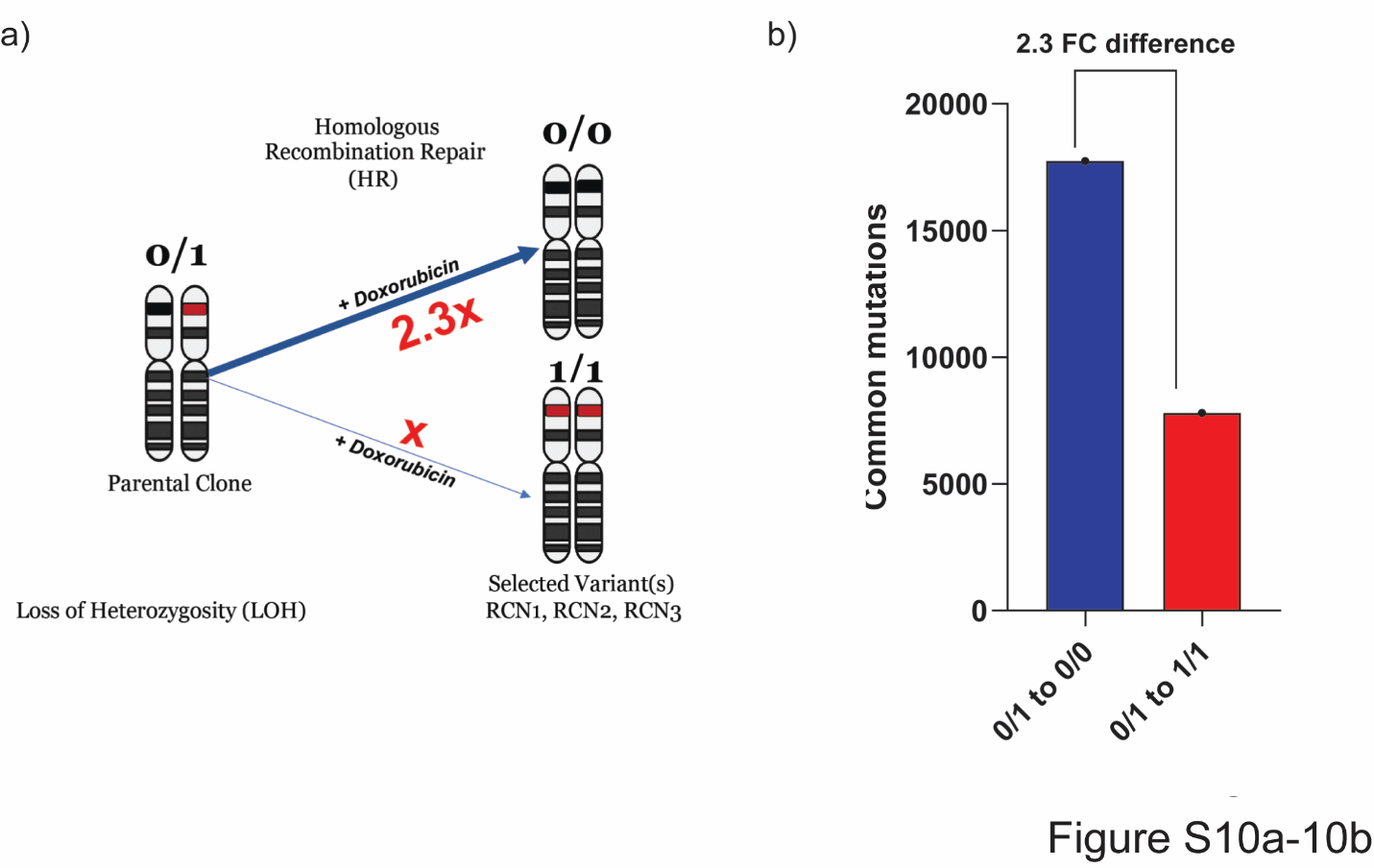

**Supplementary Figure S8**. (a) Schematic representation of allele changes associated with loss of heterozygosity (LOH) following homologous recombination (HR) repair in H1975 cells. In this context, LOH refers to the conversion of heterozygous mutated alleles to homozygous states at selected variants, such as a shift from 0/1 to 0/0 or from 0/1 to 1/1. Notably, the frequency of 0/1 to 0/0 conversions was 2.3 times higher than that of 0/1 to 1/1 in the H1975 clones. (b) The bar graph displays the number of mutations corresponding to each allele shift depicted in panel (a), illustrating the distribution of different types of allelic conversions observed in the H1975 clones.

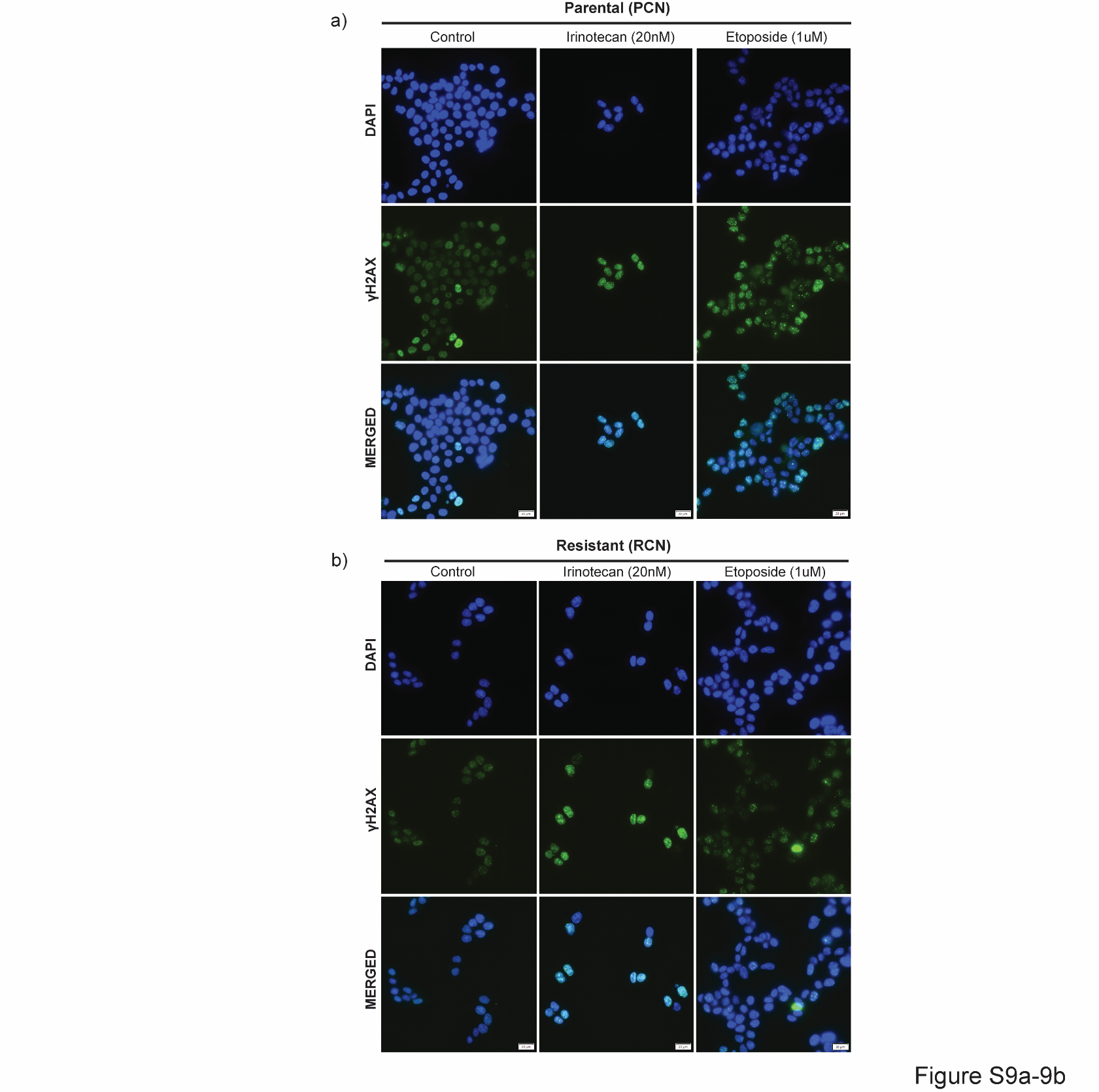

**Supplementary Figure S9.** Acquisition of doxorubicin resistance impairs the ability of etoposide but not SN-38 to induce DSBs. DNA double-strand breaks (DSBs) were assessed by monitoring γH2AX foci formation following drug treatment. Cells were exposed to etoposide (1 µM) or SN-38 (20 nM) for 1 hour, and γH2AX foci were visualized by immunofluorescence. (a) Representative images showing robust γH2AX foci formation in parental naïve cells (PCN) after treatment with etoposide or SN-38. (b) Representative images of resistant cells (RCN) under the same conditions, demonstrating reduced γH2AX foci formation following etoposide exposure, while SN-38 treatment still induces γH2AX foci. Images were analyzed with the integrated software images are at a scale of 20 µm.
