## Supplementary Figures and legends, Tables for "Systematic Generation of Mutations at Topoisomerase II Cleavage Sites Enables Cancer Adaptation to Doxorubicin Therapy": Supplemetary_file_Mutation_Resonance Probability.docx

The Mutation Resonance Problem

When a drug consistently selects for the same resistance mutations across independent populations, it indicates strong selective pressure driving convergent evolution. This suggests the drug acts through a highly specific mechanism of action, targeting a crucial protein or pathway where only a few mutations can confer resistance without compromising essential function. The limited diversity of resistance mutations reflects constrained evolutionary options and low genetic redundancy, meaning there are no alternative pathways to bypass the drug’s effect. Although mutations arise randomly, the drug acts as a powerful selective filter, allowing only rare, pre-existing or induced mutations that confer survival. In some cases, stress-induced mutagenic pathways (e.g., the bacterial SOS response) may further increase the likelihood of these adaptive mutations, giving the appearance of a directed mechanism.

The accompanying mathematical model formalizes this by estimating the **probability of overlap between two independent sets of gene responses** (A and B) within a large gene universe (U), helping to quantify how likely such convergence would occur by chance.

**Input:**

●	the genome has size of U of cardinality |U | = n

●	the number of mutations in two different clones - A and B of cardinality |A| = |B| = m

**Output:**

●	the probability P (|A ∩ B| = k) of the fact that the intersection

|A ∩ B| = k

**Main assumptions:**

*A* ⊆ *U,* *B* ⊆ *U*

|*A*| = |*B*| = *m,* |*U* | = *n*

The subsets *A* and *B* are chosen uniformly at random from all subsets of *U* of size *m*. The choices of *A* and *B* are independent.

First exact formulae:

$$\frac{Number of pairs (A, B) with |A| = |B| = m and |A \cap B| = k}{Total number of pairs (A, B) with |A| = |B| = m}$$

1. The total number of pairs *(A, B).* Clearly, the total number of pairs is:

$$\binom{n}{m} \binom{n}{m}= {\binom{n}{m}}^{2}$$

2. The number of favorable pairs. Choose *S* of size *k* for the intersection. Choose *A* to be S $\cup$ $\tilde{A}$ where $\tilde{A}$is chosen from *U - S* of size m - k. Choose *B* to be *S* $\cup$ $\tilde{B}$, where $\tilde{B}$is chosen from *U - S* of size *m - k*, but we require that $\tilde{A} \cap\tilde{B}$ = $\emptyset$ to ensure that the intersection is exactly *S*.

Hence, the number of ways to choose S is $\binom{n}{k}$.

The number of ways to choose $\tilde{A}$ is $\binom{n-k}{m-k}$

For *B*, we need to choose $\tilde{B}$ from U – S of size m – k, but disjoint from the number of ways to choose $\tilde{B}$ disjoint from $\tilde{A}$ is $\binom{n-k-(m-k)}{m-k}= \binom{n-m}{m-k}$. Therefore, the number of favorable pairs is

$$\binom{n}{k}\binom{n-k}{m-k}\binom{n-m}{m-k}$$

So the probability needed is

$$P= \frac{\binom{n}{k}\binom{n-k}{m-k}\binom{n-m}{m-k}}{{\binom{n}{m}}^{2}}$$

To simplify this expression, note that

$$\binom{n}{k}\binom{n-k}{m-k}= \frac{n!}{(n-k)!k!}\frac{(n-k)!}{(n-m)!(m-k)!}= \frac{n!}{k!(n-m)!(m-k)!}= \frac{n!}{m!(n-m)!}\frac{m!}{k!(m-k)!}= \binom{n}{m}\binom{m}{k}$$

Consequently, the probability is

$$P= \frac{\binom{n}{m}\binom{m}{k}\binom{n-m}{m-k}}{{\binom{n}{m}}^{2}}= \frac{\binom{m}{k}\binom{n-m}{m-k}}{\binom{n}{m}}$$

In summary, the probability that two randomly chosen subsets of size *m* from a universal set of size *n* have an intersection of size *k* is

$$P= \frac{\binom{m}{k}\binom{n-m}{m-k}}{\binom{n}{m}}$$

**Validity conditions:**

1. $0 \leq k \leq m \leq n,$
2. $m-k\leq n-m$, otherwise $\binom{n-m}{m-k}=0$

**Second exact formulae:**

$$\frac{\binom{m}{k}\binom{n-m}{m-k}}{\binom{n}{m}}=\frac{\frac{m!}{k!(m-k)!}\frac{(n-m)!}{(m-k)!(n-k)!}}{\frac{n!}{m!(n-m)!}}=\frac{m!^{2}(n-m)!^{2}}{(m-k)!^{2}(n-k)!n!}$$

**Some known symbolic approximations**

$$\binom{n}{k} \sim\binom{n^{k}}{k!}$$

$$n!\sim\sqrt{2\pi n}{\binom{n}{e}}^{n}$$

**Numerical approximations:**

- First calculation – for mutations present in two out of three clones

$$P=\frac{\binom{m}{k}\binom{n-m}{m-k}}{\binom{n}{m}}$$

$$n={3 \times10}^{9};m=11 \times{10}^{3};k=\sim1680$$

$P= \frac{\binom{m}{k}\binom{n-m}{m-k}}{\binom{n}{m}}= \frac{\binom{11000}{1680}\binom{{3\times10}^{9}-11000}{11000-1680}}{\binom{{3\times10}^{9}}{11000}}$ = ${1.1919 \times10}^{-7151}$

- Second calculation – for mutations present in all three clones (*m = 11000, k = 7600*):

$$n={3 \times10}^{9};m=11 \times{10}^{3};k=\sim7600$$

$$P= \frac{\binom{m}{k}\binom{n-m}{m-k}\binom{n-2m+k}{m-k}}{{\binom{n}{m}}^{2}}= \frac{\binom{11000}{7600}\binom{{3\times10}^{9}-11000}{11000-7600}\binom{{3\times10}^{9}-11000+7600}{11000-7600}}{\left( {{3\times10 \atop11000}}^{9} \right)^{2}}= 3.9795 \times{10}^{-82805}$$

**Biological conclusions:**

In the previously defined and described process examining gene responses to specific biological drugs, particularly regarding mutations, the observed patterns do not appear random.
